# Cell-type specific profiling of human entorhinal cortex at the onset of Alzheimer’s disease neuropathology

**DOI:** 10.1101/2024.12.31.630881

**Authors:** Patricia Rodriguez-Rodriguez, Wei Wang, Christina Tsagkogianni, Irena Feng, Ana Morello-Megias, Kaahini Jain, Vilma Alanko, Han-Ali Kahvecioglu, Elyas Mohammadi, Xiaofei Li, Marc Flajolet, Anna Sandebring-Matton, Silvia Maioli, Noemi Vidal, Ana Milosevic, Jean-Pierre Roussarie

## Abstract

Neurons located in layer II of the entorhinal cortex (ECII) are the primary site of pathological tau accumulation and neurodegeneration at preclinical stages of Alzheimer’s disease (AD). Exploring the alterations that underlie the early degeneration of these cells is essential to develop therapies that curb the disease before symptom onset. Here we performed cell-type specific profiling of human EC at the onset of AD neuropathology. We identify an early response to amyloid pathology by microglia and oligodendrocytes. Importantly, we provide the first insight into neuronal alterations that coincide with incipient tau pathology: the signaling pathway for Reelin, recently shown to be a major AD resilience gene is dysregulated in ECII neurons, while the secreted synaptic organizer molecules NPTX2 and CBLN4, emerging AD biomarkers, are downregulated in surrounding neurons. By uncovering the complex multicellular landscape of EC at these early AD stages, this study paves the way for detailed characterization of the mechanisms governing NFT formation and opens long-needed novel therapeutic avenues.

## Introduction

Alzheimer’s disease (AD) is characterized by a long preclinical phase, in which pathological alterations silently build up for years before the first symptoms appear. During these presymptomatic stages, amyloid beta (Aβ) pathology accumulates across the hippocampus and cortex, while neurofibrillary tangles (NFTs) of hyperphosphorylated tau protein first appear in the layer II of the entorhinal cortex (EC)^1-3^. Patients start to experience symptoms once neurodegeneration in the EC is already advanced and NFTs have spread to other brain regions^4^.

While major progress has been made to decipher the neuroinflammatory environment that accompanies neuronal pathologies, little is known about the neuronal genes and pathways that are responsible for NFT buildup in specific neurons and for their degeneration. One reason for this knowledge gap is that genome-wide association studies (GWAS) have mostly identified microglial genes until now, although whole genome sequencing studies^5-7^ and functional genomics strategies to analyze GWAS data^8^ are starting to reveal neuronal disease-associated pathways. For instance, a recent study reported that an individual with a rare gain of function mutation in the *RELN* gene (RELN COLBOS), that encodes the protein Reelin, was resilient to autosomal dominant AD, showing low tau burden in the EC despite high amyloid load, and a 25 year-delay in cognitive impairment from the expected onset^9^. Interestingly, Reelin, a secreted protein that can inhibit tau kinases via its receptors APOER2 and VLDLR, is highly enriched in ECII neurons. While a failing Reelin pathway could therefore underlie early EC pathology, evidence that this is indeed an early event in AD pathogenesis is still missing.

Uncovering the alterations that take place in the EC at the earliest stages of AD is essential to design treatment strategies that can curb clinical disease onset and progression, as therapeutic efforts might be of limited efficacy if they are implemented after a significant number of these neurons have died. Furthermore, toxic tau fibrils have been shown to spread from the EC to other regions where they can act as a template for new fibril formation^10^. The selective vulnerability of ECII also offers a unique opportunity to parse out some of the earliest molecular events in AD pathogenesis, before broader network dysfunction and inflammation perturb virtually every cell in the area and downstream pathological and compensatory mechanisms challenge the identification of pathways that initiate neurodegeneration.

Single-nucleus RNA-sequencing (snRNAseq) is a powerful tool for deciphering the molecular underpinnings of complex multicellular diseases like AD but capturing disease onset is challenging. NFT formation and degeneration of ECII neurons peak before AD symptoms become apparent^4, 11, 12^. The optimal time point to evidence changes responsible for disease onset is therefore years before a clinical diagnosis is even possible, during Braak stages I or II of NFT deposition. Difficulties in obtaining *postmortem* EC tissue at very early Braak stages has led to the study of other regions that get affected at later stages, or of the EC after a clinical diagnosis (Braak stage III or higher)^13-18^. A couple of recent single-nucleus RNA-sequencing (snRNAseq) studies included the EC from individuals at very early Braak stages^19, 20^. Unfortunately, either due to relatively low nuclei counts or to an inadequate cohort design to test incipient preclinical pathology, they fell short of revealing molecular pathways that could be responsible for neuronal pathology.

In the current study, we perform an in-depth analysis of the molecular changes taking place in the human EC at the earliest stages of AD neuropathology deposition by combining snRNAseq and fluorescence-activated nucleus sorting-sequencing (FANS-seq) for deeper profiling of neuronal populations. We compare a cohort of postmortem EC tissue from individuals devoid of any histopathological mark of AD with individuals asymptomatic at death but with both amyloid and tau pathology. This group, which would have most likely progressed into AD, represents the first stage of a biological definition of AD^21^. Accompanying a prototypical disease-associated microglia phenotype likely triggered by amyloid accumulation, we find neuronal downregulation of AD biomarkers NPTX2 and CBLN4. More remarkably, we provide compelling evidence for Reelin signaling dysregulation in ECII neurons, indicating that this pathway plays a pivotal role in the disease onset.

## Results

### Sample cohort, brain region selection and cell-type diversity identification

The earliest events that take place in the EC during the AD continuum happen before an individual meets clinical AD diagnostic criteria. Identifying these events thus requires the use of *postmortem* tissue from asymptomatic individuals with AD neuropathological hallmarks in the EC. We thus obtained *postmortem* EC samples with either a Braak II histopathological classification that we will refer to as Alzheimer’s disease neuropathology (ADN1-ADN5) or with no tau pathology (C1-C5) (**Supplementary Table 1**). We first investigated the presence of AD pathology within the EC to verify that the selected samples met our categorization criteria (absence vs. presence of AD-type neuropathology in C vs. ADN samples). Immunofluorescence staining for amyloid (6E10) and thioflavin S in the EC confirmed the absence of amyloid and tau pathology in C1-C4 and the presence of both amyloid and tau pathology in ADN2-ADN5. On the other hand, C5 and ADN1 were excluded from our comparative analysis because C5 showed extensive amyloid pathology while ADN1 had none (**Extended Data Fig. 1**). For an in-depth characterization of the different cell types of the human EC we performed single-nuclei RNA-seq (snRNA-seq) in nuclei isolated from fresh frozen EC brain tissue from C1-C5 and ADN1-ADN5. These same nuclei suspensions were also stained for neuron markers and subjected to fluorescence-activated nucleus sorting (FANS) followed by bulk RNA-seq for population-level profiling (**Fig. 1a**).

**Fig. 1:**
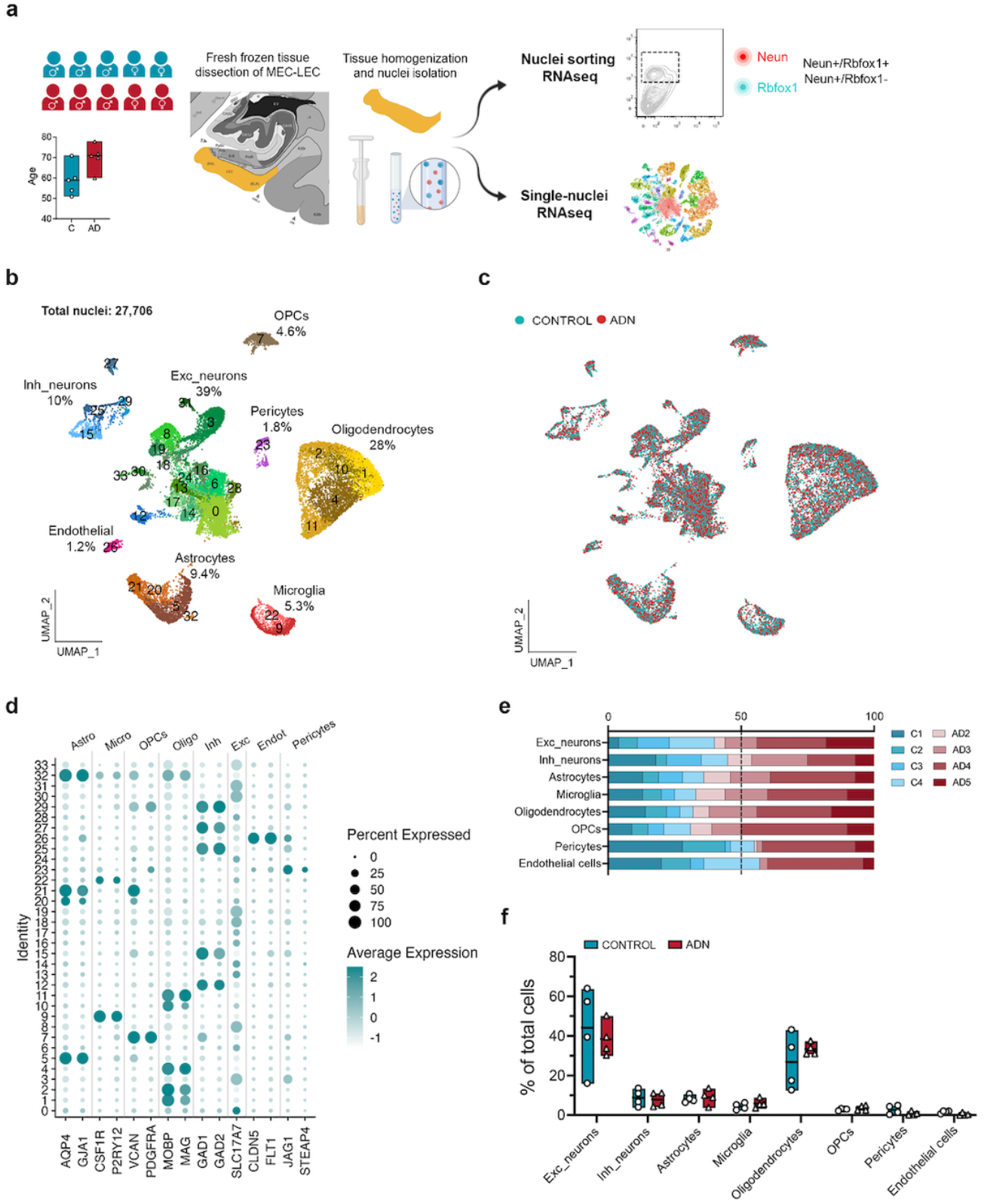
Sample cohort, experimental outline and cell type diversity identification. **a)** Schematic representation of the sample cohort and experimental approach. UMAP of the 27, 706 nuclei kept after quality control colored by b) cell-types identified (with their relative abundance) and c) by experimental group control and ADN samples. **d)** Dot plot of the expression profiles of the different cell-type signature genes. **e)** Relative contribution of the different samples to a certain cell-type population. **f)** Relative abundance of a certain cell-type population in each sample.

After filtering out low-quality nuclei (methods), we obtained expression profiles from a total of 27,706 nuclei (**Fig. 1b**). As observed in **Fig. 1c** nuclei from all samples integrated well. Clustering of these nuclei identified 8 major brain cell types based on the expression profile of literature-based cell-type specific markers^15^: 39% excitatory neurons (*SLC17A7*), 28% mature oligodendrocytes (*MOBP, MAG*), 10% inhibitory neurons (*GAD1, GAD2*), 9.4% astrocytes (*AQP4, GJA1*), 5.3% microglia (*CSF1R, P2RY12*), 4.6% oligodendrocyte progenitor cells (OPCs; *VCAN, PDGFRA*), 1.8% pericytes (*STEAP4, JAG1*) and 1.2% endothelial cells (*FLT1, CLDN5*). The percentages of each of these cell types within the total cell population are similar to previously reported snRNA-seq data from the human cortex^15^, indicating that our dataset accurately captures all major cortical cell types (**Fig. 1b and 1d**).

We first determined if the abundance of any cell type is already affected at the onset of AD neuropathology in the EC. To do so, we ran scCODA, a method that identifies compositional changes across groups (C1-C4 vs. ADN2-ADN5) for each cell type relative to the total cell population^22^. In agreement with a previous study^19^, we did not observe statistically significant differences in the abundance of any general cell type (**Fig. 1e and 1f**). We next performed pseudobulk differential gene expression (DGE) analysis for each cell-type and an in-depth analysis of the abundance and gene expression changes in their subpopulations (**Supplementary Data 1**).

### Early microglia and oligodendrocyte reactivity to amyloid pathology and potential alterations in the glymphatic system

We started our analysis by exploring the changes undergone by glial cells in the snRNA-seq data. Pseudobulk differential gene expression (DGE) followed by pathway analysis showed that, in general, glial populations are characterized by alterations in pathways related to cell stress, including changes in phagosome formation in microglia (*P* = 6.2 × 10^−3^); ubiquitination (*P* = 7.8 × 10^−5^), aggrephagy (*P* = 7.9 × 10^−4^) and heat shock stress response in astrocytes (*P* = 5.1 × 10^−5^); OPCs and oligodendrocytes also showed alterations in heat shock stress response (*P* = 1.7 × 10^−3^ and *P* = 7.1 × 10^−4^, respectively) (**Extended Data Fig. 2a-d and Supplementary Data 2**).

Studies in mice where only plaque pathology is present allow to isolate plaque-responsive gene expression programs. For instance, Chen et al. identified a set of plaque-induced genes (“PIGs”) in microglia and a myelination-related gene module (“OLIG” module) that was transiently going up in the vicinity of plaques^23^. To explore the possibility that changes in microglia and oligodendrocytes in early ADN are a direct response to plaques, we tested the overlap between the modules from Chen et al. and genes differentially expressed in ADN in microglia and oligodendrocytes. Excitingly, genes upregulated in ADN microglia and oligodendrocytes were enriched in genes from PIGs and OLIG module respectively (Wilcoxon rank sum test, *P* = 0.0041 and < 2.2 × 10^−16^ respectively), suggesting that amyloid plaques cause their upregulation (**Extended Data Fig. 2e**). Interestingly, one of the plaque-induced oligodendrocyte genes is *BIN1* (log2FC = 0.5, *P*_*adj*_ = 2.3 × 10^−03^), the second most significant genetic risk locus for late-onset AD after *APOE*^*24*^.

We then subclustered the different glial subpopulations to characterize their alterations. We did not observe any changes in cell abundance for the different OPC (OPC_s0 – OPC_s3) and oligodendrocyte subpopulations (Oligo_s0 – Oligo_s7) (**Extended Data Fig. 3a-b**). Among all the different subtypes of glial cells, one microglia subcluster, Micro_s1, was significantly more abundant in ADN samples than in controls (**Fig. 2a-b**). Since marker genes for this microglia subtype included genes characteristic of disease-associated microglia (DAM) (e.g. *TYROBP, APOE, CLEC7A, CTSZ or CD63)* (**Supplementary Data 1**), we tested if this cluster was significantly enriched in DAM signatures in mice. We found a significant enrichment of mouse DAM^25^ or “activated-response microglia”^26^ genes (Wilcoxon rank sum test *P* < 2.2 × 10^−16^ for both gene sets) (**Fig. 2c**). This cluster also significantly overlaps with a human microglia subtype previously described in *postmortem* prefrontal cortex (PFC)^14^ (MG3, Wilcoxon rank sum test *P* < 2.2 × 10^−16^) (**Fig. 2d**). To explore if amyloid plaques could trigger the Micro_s1 population, we turned again to genes identified by Chen et al. as Aβ-induced genes. These PIGs, mostly microglial, were significantly enriched exclusively in Micro_s1 (Wilcoxon rank sum test *P* = 1.2 × 10^−9^) (**Fig. 2e**), suggesting that this cluster represents microglial cells responding to amyloid accumulation.

**Fig. 2:**
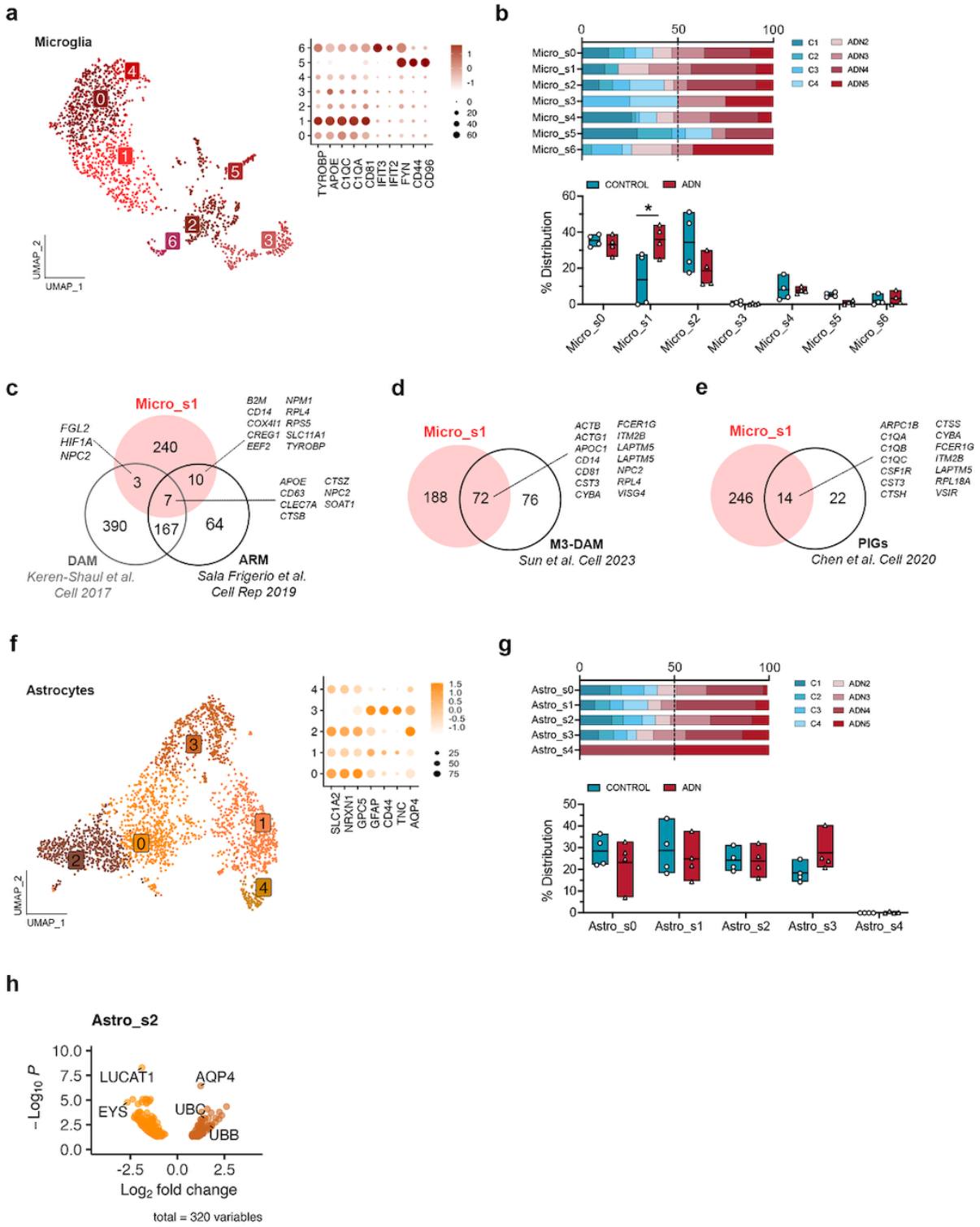
Alterations in microglia and astrocyte populations at the onset of AD neuropathology. **a)** UMAP of the different microglia subpopulations and dot plot of the expression profiles of prototypical microglia states signature genes. **b)** Upper panel: relative contribution of the different samples to a certain subpopulation; lower panel: abundance of a certain cell subtype in each sample. The asterisk represents significance in the abundance of a cell subtype by scCODA. **c)** Venn diagram of the overlap between genes enriched in Micro_s1 and DAM and ARM signature genes from 2 different studies in the mouse brain. **d)** Venn diagram showing the overlap between genes enriched in Micro_s1 and a DAM gene signature described in human AD (M3 in Sun et al.). **e)** Venn diagram of the overlap between genes enriched in Micro_s1 and plaque-induced genes (PIGs) identified in the mouse. **f)** UMAP of the different astrocyte subpopulations and dot plot of the expression profiles of prototypical astrocyte subtypes signature genes. **g)** Upper panel: relative contribution of the different samples to a certain subpopulation; lower panel: abundance of a certain cell subtype in each sample. **h)** Volcano plot of the differentially expressed genes between control and ADN in Astro_s2.

While we did not find statistically significant changes in the abundance of astrocyte subpopulations with scCODA, Astro_s3, enriched in reactive astrocyte markers (*GFAP, CD44* and *TNC)* (**Fig. 2f**), showed a trend towards increased abundance in ADN samples (**Fig. 2g**). Interestingly, however, of all astrocyte subclusters, Astro_s2, characterized by higher expression of homeostatic genes (e.g. *SLC1A2, NRXN1* and *GPC5)* (**Fig. 2h**), showed the highest number of differentially expressed genes (DEGs) by pseudobulk DGE analysis (**Supplementary Data 1**). The top upregulated gene in this subgroup of astrocytes was *AQP4* (log2FC = 1.23, *P*_*adj*_ = 3.67 × 10^−07^) (**Fig. 2e**). *AQP4* codes for the water channel aquaporin 4, essential for brain waste clearance through the glymphatic system^27-29^. This therefore suggests that changes in the function of the glymphatic system could be an early event in AD neuropathology within the EC.

### Parvalbumin and Vip interneurons dysfunction

Analysis of the DEGs in inhibitory neurons by pseudobulk DGE analysis between ADN and control samples revealed alterations in several pathways associated with intercellular communication, including integrin cell surface interactions (*P* = 6.5 × 10^−4^), tight and gap junction signaling (both pval = 0.013) and eNOS signaling (*P* = 7.9 × 10^−7^) (**Fig. 3a**) (**Supplementary Data 2**). *FBN2* was one of the top upregulated genes in these cells (log2FC = 1.3, *P*_*adj*_ = 1.5 × 10^−5^). Notably, a very recent GWAS study based on whole genome sequencing data identified a new common variant (rs147450666) proximal to the *FBN2* locus that is protective against AD^30^. These cells also showed a significant downregulation of *RELN* (log2FC = −0.66, *P* = 3.5 × 10^−2^), *SPRY4-AS1* (log2FC = −1.5, *P*_*adj*_ = 2.7 × 10^−3^) and *FGF1* (log2FC = −0.9, *P*_*adj*_ = 1.1 × 10^−2^) (**Fig. 3a and Supplementary Data 1**). *FGF1* and *SPRY4-AS1* (also known as *LOC101926941*) belong to the same genomic locus, which suggests a general downregulation of the whole locus. Interestingly, a single nucleotide polymorphism (SNP) in *SPRY4-AS1*, rs3850579, showed a suggestive association with AD in a GWAS of neuropathology-verified cases (*P* = 7.1 × 10^−6^)^31^.

**Fig. 3:**
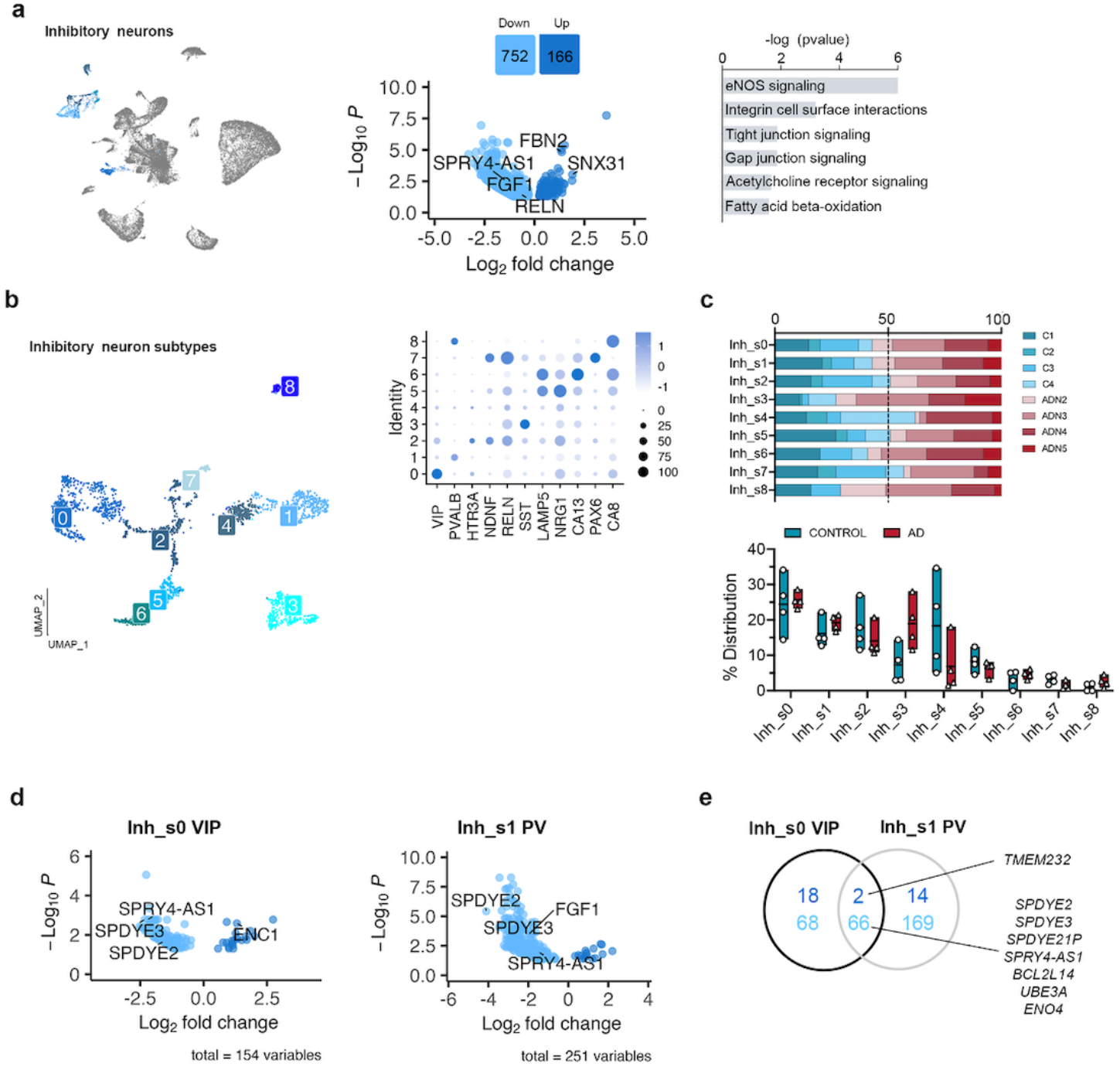
Alterations in inhibitory neuron populations at the onset of AD neuropathology. **a**) UMAP of the inhibitory neuron populations in the general clustering, volcano plot showing the differentially expressed genes between control and ADN in inhibitory neurons and dysregulated pathways in these cells. **b)** UMAP of the different inhibitory neuron subpopulations and dot plot of the expression profiles for signature genes of prototypical subtypes. **c)** Upper panel: relative contribution of the different samples to a certain subpopulation; lower panel: abundance of a certain cell subtype in each sample. **d)** Volcano plots of the differentially expressed genes in VIP and PV interneurons between control and ADN. **e)** Venn diagram showing the overlap between differentially expressed genes in VIP and PV interneurons.

Clustering of inhibitory neurons revealed 9 subclusters (Inh_s0 – Inh_s8) that represented all the major types of inhibitory neuron subpopulations (**Fig. 3b**). Analysis of the PFC at more advanced disease stages reported loss of SST and RELN-LAMP5 interneurons^13^, and a recent study in this same region on individuals with low amyloid burden found decreased levels in NDNF-PROX1 interneurons^32^. In agreement with the study by Leng et al^19^, we did not observe differences in the abundance of any interneuron subpopulation at this early stage of neuropathology deposition (**Fig. 3c**). On the other hand, pseudobulk DGE analysis showed major alterations in the transcriptional profiles of VIP (Inh_s0) and parvalbumin (PV, Inh_s1) interneurons (**Fig. 3d and Supplementary Data1**). Genes from the Speedy/RINGO family (*SPDYE2, SPDYE3 –*located in a genome-wide significant AD locus^33^- and *SPDYE21P*) and regulators of cyclin-dependent kinases were commonly downregulated in both interneuron subtypes (**Fig. 3e**). Other gene expression changes were subtype-specific, like *ENC1* upregulation in VIP interneurons (log2FC = 1.31, *P*_*adj*_ = 2.6 × 10^−3^), an autophagy modulator that has been suggested to play a role in cognitive resilience in the presence of ageing-related neuropathologies^34^. *FGF1* was downregulated in PV interneurons (log2FC = −1.97, *P*_*adj*_ = 3.3 × 10^−4^). Together these results suggest the presence of early abnormalities in inhibitory neurons, in particular PV and VIP inhibitory neurons.

### Excitatory neuron subpopulations in the human entorhinal cortex and their alterations at early AD neuropathology

Pseudobulk DGE and pathway analysis of EC excitatory neurons showed alterations in CREB (*P* = 0.003), nNOS (*P* =0.03) and HIFα (*P* = 0.028) signaling (**Fig. 4a and Supplementary Data 2**). Changes in these pathways are suggestive of early alterations in neuronal excitability that can accompany tau accumulation^35^. Interestingly, *RELN* was highly downregulated in these cells (log2FC = −2.88, *P*_*adj*_ = 4.2 × 10^−4^) (**Fig. 4a and Supplementary Data 1**).

**Fig. 4:**
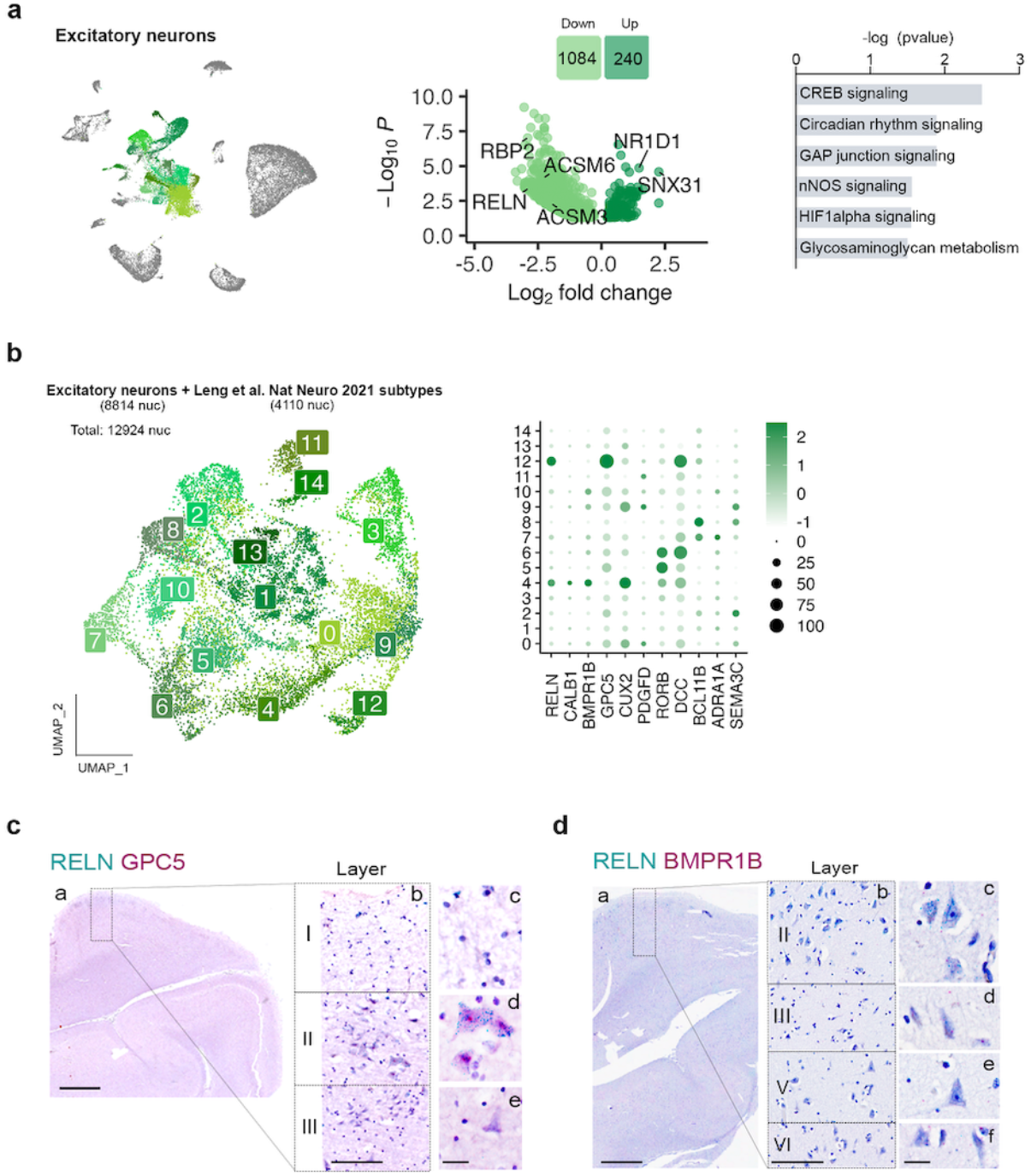
Gene expression changes in excitatory neurons and subpopulation characterization. **a)** UMAP of the excitatory neuron populations in the general clustering, volcano plot showing the differentially expressed genes between control and ADN in excitatory neurons and dysregulated pathways in these cells. **b)** UMAP of the different excitatory neuron subpopulations in the integration of our dataset with Leng et al. and dot plot of the expression profiles of layer-specific subtype markers. **c)** In-situ hybridization images of RELN (green) and GPC5 (red) in the human EC. Scale bar 2 mm (a), 250 μm (b) and 20 μm (c-e). **d)** In-situ hybridization images of RELN (blue) and BMPR1B (red) in the human EC. Scale bar 2 mm (a), 250 μm (b) and 20 μm (c-f). In situ hybridization are counterstained with hematoxylin.

The large diversity of excitatory neuron subpopulations can result in overall modest numbers of nuclei for each of them, making it difficult to accurately identify cell subtypes. To overcome this limitation, we integrated excitatory neurons from a previously published dataset on the human EC at very early Braak stages (Leng et al.^19^; Braak 0 or II, 3 and 4 samples respectively) with our excitatory neurons (**Fig. 4b**). As observed in **Extended Data Fig. 4a**, samples from both datasets integrated well.

This approach allowed us to identify a total of 15 excitatory neuron subpopulations (Exc_s0 – Exc_s14) (**Fig. 4b**). Although molecular markers are still missing for the various neuron subtypes of human EC, *RELN* is known to be expressed at high levels in perforant path-forming layer II neurons (stellate/fan) across species, including humans^36-38^, in addition to several inhibitory neuron subpopulations^36-38^. Among our excitatory neuron populations, Exc_s4 and Exc_s12 were significantly enriched in *RELN* (**Supplementary Data 1**). These clusters might therefore constitute vulnerable layer II neurons. Besides *RELN*, Exc_s4 was also enriched in *CALB1*, a marker for pyramidal neurons from layer II/III in non-human primates and mouse^39, 40^, and in *BMPR1B*, previously suggested to be enriched in layer II neurons^41^. Exc_s12, on the other hand, was enriched in *GPC5*. To validate the identity of these populations, we performed in situ hybridization (ISH) on formalin-fixed, paraffin-embedded sections from the human EC of control cases (C9-C10). As shown in **Fig. 4c**, neurons stained for both *RELN* and *GPC5* were large neurons of polygonal shape in layer II, displaying dark hematoxylin counterstain and forming islands^42^, consistently reported as early sites of NFT deposition. Cells stained for *RELN* and *BMPR1B* tended to have pyramidal shape and to be located in layer III, consistent with their expression of *CALB1*^*37*^ (**Fig. 4d)**.

We did not find a significant difference in cell abundance for any of the excitatory neuron subpopulations, including layer II neurons (**Extended Data Fig. 4b)**. Leng et al. described a significant loss of a *RORB*-enriched population of neurons from a superficial layer of the EC at Braak II. In our integrated data, we indeed found a decrease in *RORB* enriched Exc_s5 and Exc_s6 nuclei at Braak II vs 0 in the Leng et al. dataset, while our Braak 0 and II samples showed similar abundance of these populations (**Extended Data Fig. 4c**). To investigate the location of *RORB*-expressing neurons in our human EC sections, we performed duplex ISH for *RELN* and *RORB* and did not find significant colocalization between the two markers. *RORB*-expressing neurons were mostly layer V pyramidal neurons (**Extended Data Fig. 4d)**.

Pseudobulk DGE analysis between C1-C4 and ADN2-ADN5 for the different excitatory neuron subpopulations did not find significant DEGs for most subpopulations, likely because of their low abundance.

### Transcriptional changes in EC neurons at the onset of AD neuropathology

To further understand molecular changes in EC neurons, we took advantage of the deeper molecular profiling allowed by fluorescence-activated nucleus sorting-sequencing (FANS-seq). NeuN is a nuclear antigen present in all neurons. Another nuclear antigen, RBFOX1 (Fox-1 homolog A), is enriched in inhibitory neurons derived from the medial ganglionic eminence (MGE), such as SST and PV interneurons, but relatively impoverished in most interneurons originating from the caudal ganglionic eminence (CGE)^43^, offering a way to gain in cell-type resolution by sorting nuclei based on RBFOX1 and NeuN levels. Indeed, our snRNAseq data showed higher *RBFOX1* expression in Inh_s1 (PV), Inh_s3 (SST) and Inh_s8 (PV-CA8). Inh_s5 (LAMP5-NRG1) and Inh_s6 (LAMP5-CA13) were also characterized by high *RBFOX1* expression levels (**Extended Data Fig. 5b**). Little is known about RBFOX1 levels across different excitatory neuron types. RBFOX1 immunostaining on EC sections showed a range of nuclear RBFOX1 across EC layers: low nuclear RBFOX1 in layer III excitatory neurons, high levels in layer V excitatory neurons and intermediate levels in layer II RELN+ excitatory neurons (**Extended Data Fig. 5c**). We thus sorted neuron nuclei (C1-C4 and ADN2-ADN5) using the pan-neuronal marker NeuN and separated them based on RBFOX1 levels into neurons positive for both NeuN and RBFOX1 (NeuN^+^/RBFOX1^+^), i.e. layer V excitatory and MGE-derived interneurons; or NeuN positive and RBFOX1 negative neurons (NeuN^+^/RBFOX1^−^), i.e. layer III excitatory and CGE-derived interneurons (**Extended Data Fig. 5a**) and performed bulk RNA-seq in the 2 populations.

As expected, *LHX6* and *SP9*, transcription factors necessary for the specification of MGE-derived interneurons^44-48^, were enriched in NeuN^+^/RBFOX1^+^ nuclei (log2FC = 3.24, *P*_*adj*_ = 2.3 × 10^−12^ and log2FC = 1.70, *P*_*adj*_ = 0.031 respectively). *SST* and *PVALB* were also among the top enriched genes in this population (log2FC = 3.07, *P*_*adj*_ = 3.2 × 10^−3^ and log2FC = 2.94, *P*_*adj*_ = 1.6 × 10^−4^, respectively). On the other hand, the NeuN^+^/RBFOX1^−^ population was enriched for *SP8* (log2FC = 2.08, *P*_*adj*_ = 0.015) and *NR2F2* (log2FC = 1.79, *P*_*adj*_ = 1.1 × 10^−4^), transcription factors necessary for CGE interneuron specification^49-51^ (**Extended Data Fig. 5b and Supplementary Data 3**). However, these cells were not enriched in prototypical markers of interneurons derived from the CGE, like *VIP* or *NDNF*, indicating that RBFOX1 might be impoverished only in some subsets of these populations. Instead, we found an enrichment in markers of more specific subsets of CGE-derived interneurons like *CHRNA7* (log2FC = −2.0, *P*_*adj*_ = 4.8 × 10^−15^), a marker for a subset of NDNF-positive neurons from layer I^52^ and *CALB2* (log2FC = −2.10, *P*_*adj*_ = 2.0 × 10^−4^), a marker for a subset of VIP-positive interneurons^49^ (**Extended Data Fig. 5b**). Regarding excitatory neurons, in agreement with nuclear RBFOX1 levels, layer V neurons were enriched in the NeuN^+^/RBFOX1^+^ and layer II/III neurons in the NeuN^+^/RBFOX1^−^ population. Layer II RELN+ excitatory neurons seemed to be distributed within both populations (**Extended Data Figs. 5d-e**).

The NeuN^+^/RBFOX1^−^ population showed 10 DEGs in ADN compared to control (**Supplementary Data 3**). Strikingly, the only two downregulated genes, *NPTX2* (log2FC = −1.79, *P*_*adj*_ = 0.0011) and *CBLN4* (log2FC = −2.34, *P*_*adj*_ = 0.037), are important regulators of the excitation/inhibition (E/I) balance by promoting the maturation of synapses^53^, and two prominent early downregulated AD biomarkers^54, 55^ (**Fig. 5a**). Our data suggests that this decrease occurs transcriptionally in a subset of EC neurons. To identify the neuron type responsible for this effect we investigated the expression of *NPTX2* and different markers of CGE-derived interneurons – *HTR3A, VIP* and *PAX6* – in control EC by ISH, and found very little co-expression (**Extended Data Fig. 5f**). The bulk of *NPTX2* signal was instead found in cells with pyramidal morphology and larger nuclei. Interestingly, according to our snRNAseq data, *NPTX2* is enriched in two subpopulations of excitatory neurons: Exc_s0 and Exc_s9, both sorted in the NeuN^+^/RBFOX1^−^ fraction and enriched in *LAMP5* (**Extended Data Fig. 5d**). Using multiplexed ISH, we indeed found that 60.8% of *LAMP5* positive layer II/III neurons also expressed *NPTX2* (**Extended Data Fig. 5f and 5g**). We also observed *NPTX2* in layer V (**Extended Data Fig. 5g**), but, as mentioned before, these neurons are not enriched in the NeuN^+^/RBFOX1^−^ fraction and thus cannot be responsible for its downregulation. Interestingly, *CBLN4* is also enriched in Exc_s9 neurons in our snRNAseq dataset (log2FC = 1.44, *P*_*adj*_ = 0.0089). Indeed, we also find it to be enriched in *LAMP5* positive layer II/III neurons by ISH (62 % of all *LAMP5* positive neurons), with expression in *LAMP5* negative layer II neurons as well (**Extended Data Fig. 5g**). While different neuron populations could independently downregulate *NPTX2* and *CBLN4*, it is noteworthy that are co-expressed in 42% of *LAMP5* positive layer II/III pyramidal neurons (**Fig. 5b and Extended Data Fig. 5f**). This neuron population could therefore be uniquely affected in early ADN and downregulate both genes.

**Fig. 5:**
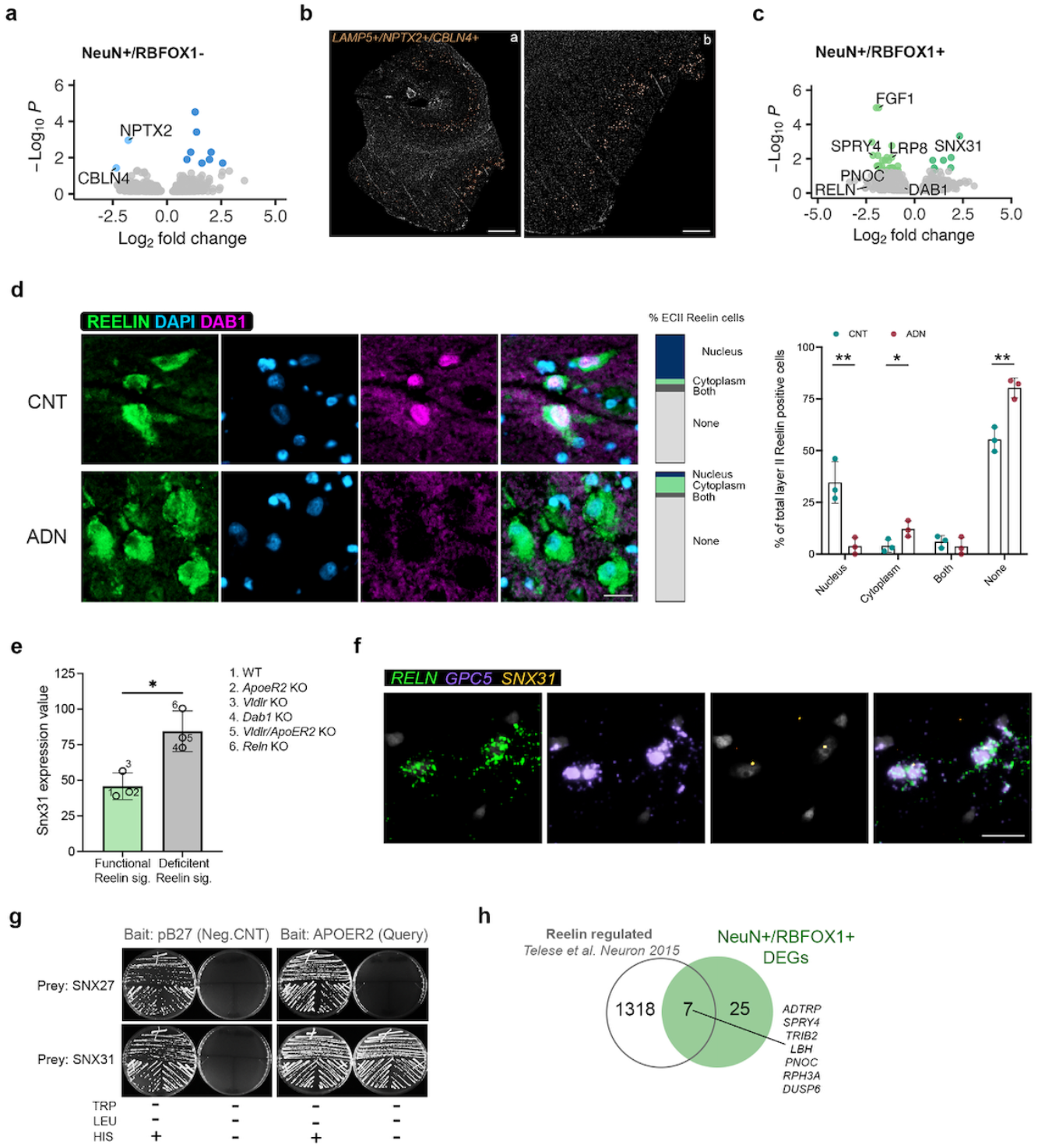
Early alterations in Reelin signaling in EC neurons. **a)** Volcano plot of the differentially expressed genes in the NeuN^+^/RBFOX1^−^ population. **b)** ISH images showing the spatial localization of LAMP5+/NPTX2+/CBLN4+ cells in the human EC. Scale bar 2 mm (a) and 800 μm (b). **c)** Volcano plot of the differentially expressed genes in the NeuN^+^/RBFOX1^+^ population. **d)** Confocal microscopy images of Reelin (green) and Dab1 (pink) in the EC of control (CNT) and ADN samples. The bars and the column graph represent the subcellular localization of Dab1 in % of total Reelin positive cells quantified. Nuclei were stained with DAPI (blue). Scale bar 50 μm. **e)** Snx31 expression levels in different mouse models either can have functional reelin signaling (WT, Vldlr KO and ApoER2 KO, green bar) or that have deficient reelin signaling (Vldlr/ApoER2 KO, Dab1 KO and Reln KO, grey). Re-analysis of GEO dataset GSE94896. **f)** ISH images of RELN, GPC5 and SNX31 localization in the human EC. Scale bar 20 μm. **g)** Yeast one-by-one solid grow test for SNX27 and SNX31 binding to the intracellular domain of APOER2. **h)** Venn diagram showing the overlap between Reelin regulated genes and differentially expressed genes in NeuN^+^/RBFOX1^+^ neurons.

DGE analysis between control and early AD neuropathology in the NeuN^+^/RBFOX1^+^ population showed a total of 32 DEGs (**Supplementary Data 3**). The top downregulated gene was *FGF1*, which corroborated the decrease seen in the PV interneurons by snRNAseq. Strikingly, of the few DEGs, *LRP8*, encoding ApoER2, a receptor for both the main AD genetic susceptibility factor APOE and for Reelin, was significantly downregulated in ADN brains (log2FC = −1.19, *P*_*adj*_ = 8.5 × 10^−3^). This was accompanied by a nominally significant downregulation of both *RELN* (log2FC = −2.42, *P* = 9.5 × 10^−3^) and *DAB1* (log2FC = −0.54, *P* = 1.5 × 10^−2^), the main transducer of the Reelin signaling pathway (**Fig. 5c**). Interestingly, a previous GWAS study in APOEe4 homozygous carriers identified a significant association of the *DAB1* locus with AD onset^56^. To explore potential alterations in Reelin signaling, we performed immunofluorescence staining for Reelin and Dab1 in fresh-frozen tissue sections from a different sample cohort (C6-C8 and ADN6-ADN8, **Supplementary Table 1**). Dab1 was only detectable in a fraction of Reelin positive ECII neurons, potentially owing to its degradation upon Reelin receptors activation^57, 58^. In cells where Dab1 was detectable, it could either be found in the cytoplasm or in the nucleus, confirming previous reports of the nuclear-cytoplasmic shuttling of the protein^59, 60^. Within ECII Reelin-positive cells, a higher percentage of cells showed positive Dab1 signal in control samples than in ADN, as shown in **Fig. 5d**. Interestingly, when present, Dab1 was preferentially located in the nucleus in control conditions and in the cytoplasm in ADN samples. It is likely that cytoplasmic Dab1 is found in cells with inactive Reelin signaling, suggesting impaired signaling in ADN.

Among the 7 upregulated genes in NeuN^+^/RBFOX1^+^ neurons, *SNX31* was the most significantly changed (log2FC = 2.33, *P*_*adj*_ = 4.72 × 10^−4^) (**Fig. 5c**). *SNX31* was also among the top upregulated genes in the pseudobulk DGE analysis of both inhibitory (log2FC = 1.89, *P*_*adj*_ = 9.3 × 10^−4^) and excitatory neurons (log2FC = 2.25, *P*_*adj*_ = 2.6 × 10^−5^) (**Fig. 3a and 4a**). Interestingly, *SNX31* was previously shown to be upregulated in neurons in ApoER2 KO mice and strongly downregulated when rescuing these neurons with ApoER2 intracellular domain (ICD), suggesting an ApoER2-ICD-mediated repression of its transcription^61^. To further explore the association between Reelin signaling and *SNX31* expression we explored a publicly available dataset of microarray gene expression analysis in WT and Knock-out (KO) mice for *Vldlr, ApoER2*, double *Vldlr/ApoER2* KO, *Dab1* and *Reln* (Dillon et al., Neuron 2017, Gene expression omnibus accession GSE94896)^62^. Excitingly, their results showed a clear upregulation of *Snx31* in all the models with impaired Reelin signaling (*Dab1* KO, *Reln* KO and double *Vldlr/ApoER2* KO) (**Fig. 5e**).

The precise role of the FERM-domain containing sorting nexin encoded by *SNX31* is unknown. However, its closest paralogue, SNX17 (39% amino acid sequence homology, also containing a FERM domain) was identified by a yeast two-hybrid screen as a binding partner of the ApoER2 intracellular domain and demonstrated to activate Reelin signaling by promoting ApoER2 recycling^63, 64^. SNX31 might thus also be a regulator of the Reelin signaling pathway through similar mechanisms. To investigate this possibility, we tested the interaction between SNX31 and the cytoplasmic domain of ApoER2, using a yeast 1-by-1 test – a targeted version of the yeast two-hybrid method used to identify the direct interaction between SNX17 and APOER2^64^. We found a direct interaction between the cytoplasmic domain of ApoER2 and SNX31 (likely via the NPxY motif) but not SNX27, another FERM-domain containing sorting nexin (**Fig. 5g**), suggesting that, like SNX17, SNX31 could regulate Reelin signaling. SNX31 is expressed at very low levels in control conditions, but we could detect its expression at Braak stage II in RELN+ GPC5+ ECII neurons using ISH (**Fig. 5f**), suggesting that ECII neurons might indeed use SNX31 as a mechanism to compensate for decreased Reelin signaling.

One of the outcomes of a functional Reelin pathway is ApoER2 ICD-mediated transcription. Impaired Reelin signaling would thus cause dysregulation of ApoER2 transcriptional target genes. Indeed, we compared our DGE data with a published set of Reelin/ApoER2 regulated genes^65^ and found that genes transcriptionally induced by Reelin were significantly downregulated in ADN samples in both neuronal subfractions (Wilcoxon rank sum test *P* = 1.04 × 10^−4^ in NeuN^+^/RBFOX1^+^ and 1.31 × 10^−5^ in NeuN^+^/RBFOX1^−^ neurons) (**Fig. 5h**). Altogether, our data suggest a robust dysregulation of the Reelin signaling pathway in the EC, both in vulnerable layer II neurons and in neighboring neurons (**Extended Data Fig. 6**).

## Discussion

In the current study we sought to capture the transcriptional events associated with incipient NFT formation during the silent phase of AD, this is, when NFT pathology is still largely confined to the EC. To do so, we analyzed postmortem EC from cases with either no tau (Braak 0) or amyloid pathology in the EC or with both amyloid and early tau pathology (Braak II).

A unique aspect of this study is the combination of FANS-seq and snRNAseq to identify alterations in neuron subpopulations. Differential gene expression analysis from snRNAseq data notoriously suffers from gene expression variability and high dropout rate. On the other hand, while the cell type resolution of FANS-seq is often limited by the availability of appropriate nuclear markers, it provides deep sequencing, detection of lower expressed genes and yields robust differential expression analyses between groups^66^. This approach allowed us to identify a significant decrease in *LRP8*, not found by snRNAseq, in one of our neuronal fractions. This, together with nominally significant decreases in *DAB1* and *RELN* in these same neurons and with a significant downregulation of *RELN* in the pseudobulk DGE analysis of both excitatory and inhibitory neurons, prompted us to explore possible alterations in Reelin signaling. Strikingly, immunofluorescence analysis confirmed impaired Reelin signaling in ECII neurons: Dab1was detectable in a greater fraction of ECII neurons in control than ADN samples and, when detectable, was preferentially nuclear in control vs. cytoplasmic in ADN. It is likely that Dab1 is transported to the nucleus by piggybacking on the ApoER2 ICD, which is cleaved upon Reelin binding and shuttles to the nucleus to regulate gene transcription. In the nucleus, Dab1 has been shown to enhance ApoER2 ICD mediated transcription^67^. The nuclear absence of Dab1 in ADN would result in the downregulation of Reelin transcriptional targets, which we indeed observed in both sorted neuron populations. Additional evidence of a waning Reelin signaling comes from the upregulation of *SNX31*, which is repressed by Reelin/ApoER2 ICD signaling, and we further show is an interaction partner of the ApoER2 ICD. Like its paralogue SNX17, it could therefore regulate ApoER2 recycling to the cell surface^63^. This would make it an important negative feedback modulator of Reelin signaling and, together with its recent association with AD in a genetic study in individuals of African ancestry^68^, a compelling therapeutic target to counteract these early pathological changes.

According to our data, a few non-mutually exclusive mechanisms could contribute to Reelin signaling alterations: 1) they could be due to reduced Reelin synthesis or secretion by *RELN*-expressing neurons in the EC. While this is supported by our pseudobulk DGE analysis and by previous studies^38^, we still detect enrichment for Reelin in ECII neurons by immunofluorescence in ADN samples comparable to controls. Nevertheless, we cannot rule out a loss of Reelin in subsets of ECII neurons; 2) heparan sulfate proteoglycans might be different in ADN. They stabilize Reelin in the extracellular matrix^69^, and their increased interaction with the *RELN* COLBOS variant might be responsible for its protective effect^9^. It is noteworthy that *GPC5*, which codes for a heparan sulfate proteoglycan highly enriched in ECII neurons, showed a nominally significant downregulation in our sorted neurons (log2FC = −2.48, *P* = 0.013). 3) Reelin receptor levels or function might be decreased. This is supported by the significant downregulation of *LRP8* and the suggestive decrease in *DAB1* expression in our dataset. Evidence for this downregulation was also found in a previous study at later disease stages^70^. Furthermore, Reelin shares its receptors VLDR and ApoER2 with apolipoprotein E (ApoE), the main genetic risk for late-onset AD. Both proteins compete for receptor binding^71^, and ApoE modulates the recycling of ApoER2 to the cell surface^72^. DAM microglia, which showed higher abundance in our ADN samples, likely in the vicinity of plaques, strongly upregulate *APOE*, which is also a prototypical DAM marker^25, 26^. The ApoE-enriched microenvironment around plaques might thus affect Reelin signaling in nearby neurons through changes in ApoER2 cell surface availability.

Impairment in Reelin signaling could have consequences on neighboring neurons and might be the cause of the downregulation of *NPTX2*, a transcriptional target of Reelin^67^ in one of our sorted neuron populations. Decreased NPTX2 protein level in CSF is a very consistent feature of early AD stages^73, 74^ and correlates with cognitive decline, being a more sensitive CSF biomarker than tau or p-tau^75^. Furthermore, CBLN4 has also been found to be the top downregulated protein in CSF in tau positive vs. tau negative amyloid positive patients^76^. A parsimonious explanation for *NPTX2* and *CBLN4* downregulation in the NeuN^+^/RBFOX1^−^ population would be that a common neuron type particularly affected in early ADN downregulates both. We found co-expression of these genes in an RBFOX1^−^ pyramidal neuron population in layer II/III mainly located in the lateral edge of the EC, area 35a, known to be the first to present NFTs^77^. Dysregulation of these genes could have extensive consequences on the excitation/inhibition balance of the region.

Interestingly, we also observed extensive gene expression changes in PV and VIP interneurons in ADN compared to control, including a downregulation of *FGF1* in PV interneurons that we also found in one of our sorted neuron populations. PV interneurons are in close contact with layer II excitatory neurons in the lateral and medial entorhinal cortex, where they play an essential role in feedback inhibition^78, 79^. Furthermore, they have been previously reported to undergo morphological alterations in the ECII at early AD stages^80^. These results indicate that alterations in the function of PV interneurons may contribute to widespread alterations in the excitation/inhibition balance of the superficial layers of the EC.

We also evidenced transcriptional alterations in glial cells. By comparing genes induced in ADN in oligodendrocytes and markers enriched in the Micro_s1 microglia subpopulation with previous sets of amyloid-plaque-induced genes, we found that these cells are responding to the presence of amyloid plaques in their vicinity. Even though the DAM microglia subtype, which significantly overlaps with our Micro_s1, has been abundantly described in mouse models of amyloid pathology, they have not been clearly observed in human AD. The use of previous-generation sequencing reagents by Leng et al.^19^, Mathys et al.^15^ and Grubman et al.^18^ possibly precluded accurate capture of microglial transcriptomes. However, in a more recent study in the PFC, the microglia cluster most similar to ours and enriched in mouse DAM markers, MG3, was not more abundant in AD^20^. Whether this microglia population is specific to the EC or to early disease stages when microglia cells are purely responding to amyloid pathology will need to be explored in greater detail. One of the ADN-induced genes in oligodendrocytes also found to be upregulated by amyloid plaques is *BIN1*, located in one of the major genetic risk loci for AD. *BIN1* was linked to decreased EC thickness in AD patients^81^, as well as with higher tau-PET burden in cognitively normal individuals^82^. This early upregulation in *BIN1* may therefore contribute to AD pathology at early stages.

By focusing on the earliest pathology in the most vulnerable region to AD, this study provides critical insights into one of the field’s most enigmatic question, i.e. what causes the early degeneration of specific neuron subpopulations? Indeed, the main pathway that we find to be impaired in ECII neurons is also one of their most salient characteristics, since they are among the rare excitatory neurons that express high levels of *RELN* in the adult brain. We hypothesize that, for reasons that remain to be understood, the normal function of ECII neurons puts them at risk of forming pathological tau lesions, unless they are protected by the strong anti-tau phosphorylation and neuroprotective effects of Reelin. The notion that Reelin signaling is a gatekeeper for the initiation of tau pathology in the EC is further reinforced by the protective *RELN* COLBOS mutation and by a previous study showing that ApoER2 could be enriched in regions that develop tau pathology early in the disease continuum, including the EC^70^. Alternatively, ECII neurons might be highly dependent on Reelin signaling for other reasons than mere protection against tau hyperphosphorylation. Regardless, in the context of ADN, decreased Reelin signaling in ECII neurons would activate the kinase GSK3β^83^, involved in tau phosphorylation, while also causing regional excitation/inhibition imbalance through changes in neighboring neurons. Further research is needed to elucidate how various AD risk factors affect this axis and the interplay between Reelin and ApoE in the EC. However, our demonstration that the pathway is involved in the time and place where tau pathology originates reveals an unprecedented therapeutic opportunity to halt the disease before irreversible damage occurs in key memory-forming regions.

## Methods

### Postmortem human brain samples

A total of 5 control (C1-C5, Braak 0) and 5 early AD neuropathology (ADN1-ADN5, Braak II) fresh frozen EC brain tissue samples were provided by the Bellvitge Hospital Brain Bank (Barcelona, Spain). Additionally, paraffin-embedded sections were taken from the immediately adjacent posterior (caudal) region of these same samples. For immunofluorescence and RNAscope validation of early pathological alterations and layer II-specific excitatory neuron identification, we further obtained fresh frozen tissue slides from the hippocampal formation, including the EC, of 3 control (C6-C8, Braak 0) and 3 early AD neuropathology (ADN6-ADN8, Braak I-II) from The Netherlands Brain Bank (Netherlands). Sample information is detailed in **Supplementary Table 1**.

### Single-nuclei isolation from postmortem frozen brain tissue

Nuclei from C1-C3 and AD1-AD3 were isolated using the following protocol: Tissue was placed into a glass homogenizer containing 4 ml of cold lysis buffer (20 mM HEPES-KOH, 5 mM MgCl2, 150 mM KCl, 0.5 mM DTT, protease inhibitors, 0.2 U/ml RNAsin, 0.1 U/ml SUPERase-In and 5 μg/ml actinomycin D) and homogenized using a motor-driven Teflon-glass homogenizer (900 rpm). Homogenates were then transferred to non-stick microcentrifuge tubes and pelleted by centrifugation (1200 x g, 10minutes, 4°C). Pelleted nuclei were resuspended in 2 ml of 29% iodixanol at 4°C and centrifuged 2 times at 10,000 xg for 10 min. Between each centrifugation the upper layer formed, composed of myelin, was removed using a tissue paper coated pipette tip. After the last centrifugation, the layer corresponding to the nuclei was transferred to a new tube and nuclei were pelleted by centrifugation (10,000 x g, 10 min at 4°C). Nuclei were then resuspended in 100 μl of resuspension buffer (1% RNAse free BSA in PBS supplemented with 5μg/ml actinomycin D, SUPERase-In, RNAsin and 10 μM Vybrant DyeCycle Ruby (#V10273, ThermoFisher) stain). Single nuclei were sorted based on DyeCycle Ruby signal and submitted for sequencing in a Chromium 10x genomics setup at the Rockefeller University genomics resource center.

Because of clogging problems in the Chromium X, we had to slightly modify our protocol by loading our samples without prior DyeCycle-Ruby-based singlet sorting for C4-C5-AD4-AD5. As we had encountered problems with 2 clogged samples (AD2 and AD3), we run new samples from these same 2 individuals but performing nucleus isolation following the 10x recommended protocol (of note, the initial nucleus preparation was used for FANS-seq so that all samples underwent the same preparation for that part of our study). Briefly, tissue was lysed into a glass homogenizer containing 2 ml of cold lysis buffer (10 nM Tris-HCl, 3 mM MgCl2, 10 mM NaCl, 0.5 mM DTT, protease inhibitors, 0.2 U/ml RNAsin, 0.1 U/ml SUPERase-In and 5 μg/ml actinomycin D). They were then run through a MACS SmartStrainer into a new tube and pelleted by centrifugation (500 xg, 5 min, 4°C). Pelleted nuclei were washed twice in Nuclei Wash Buffer (1% BSA in PBS supplemented with RNAsin and SUPERase-In) and incubated for 15 min with Myelin Removal Beads. Nuclei were once more washed in nuclei wash buffer and run through an LS column. Nuclei were then separated in a sucrose gradient, pelleted and resuspended in nuclei wash buffer prior to being sequenced in a Chromium X setup. All single-nucleus RNAseq libraries were constructed using the 10X 3’ single-cell RNA-seq v3 kit following the manufacturer’s recommendations.

### Nuclei labeling and sorting

Isolated nuclei were stained following a protocol adapted from Xu, et al. *Elife* 2018^66^. Briefly, they were incubated for 20 min at 4°C in staining buffer (1% RNAse free BSA in PBS), supplemented with RNAse inhibitors (40U/ml RNAsin, #N2515, Promega and 20U/ml Superasin, #AM2696, ThermoFisher). The antibodies used were NeuN-Alexa405 (1:3,000, #NBP1-92693AF405, Novus Biologicals) and RBFOX1-Alexa594 (1:5,000, #862706, Biolegend), respectively. Nuclei were then washed with 500 μl of staining buffer and spined down at 2000 xg, 5 min. Nuclei were finally resuspended in 100 μl of resuspension buffer (1% RNAse free BSA in PBS supplemented with 5μg/ml actinomycin D, Superasin, RNAsin and 10 μM Vybrant DyeCycle Ruby (#V10273, ThermoFisher) stain) and sorted with a FACSAria cell sorter with an 85 μm nozzle. The gating strategy for sorting is depicted in **Extended Data Fig. 5A**. Briefly we first selected single nuclei based on DyeCycle Ruby staining and then we sorted NeuN^+^/RBFOX1^+^ and NeuN^+^/RBFOX1^−^ neuronal populations.

### RNA-sequencing

Pooled sorted nuclei for each neuron subpopulation were lysed in the lysis buffer provided by the RNeasy Plus Micro RNA extraction kit (#74034, Qiagen) - completing sorted nuclei up to 350 μl of RLT buffer supplemented with β-mercaptoethanol. RNA extraction was performed according to the manufacturer’s instructions, including column-based genomic DNA elimination, eluting RNA purification columns with 20 μl of RNAse free water. The volume of the RNA samples was then reduced to 5 μl using a SpeedVac concentrator at maximum speed. Library preparation was then performed with the entirety of this RNA sample, using the Trio RNA-seq kit (Tecan). The number of PCR cycles used for the amplification of the libraries was optimized using qPCR with Eva-Green as recommended in the kit. The obtained libraries were then sequenced with an Illumina NovaSeq sequencer with an SP flowcell, 50 base pairs, paired-end, at the Genomics Resource Center of The Rockefeller University.

### Differential gene expression analysis from sorted nuclei

After sequencing, adapter and low-quality bases were trimmed from the raw sequencing files in FASTQ format by fastp^84^. Reads were then aligned to the hg38 reference genome with STAR version 2.7.1a^85^. The Reads Per Kilobase of transcript per Million mapped reads (RPKM) for all genes in each sample were calculated with the R package edgeR^86^. To analyze differential gene expression between samples, DESeq2^87^ was used, applying the standard comparison mode between two experimental groups. Enrichment of different gene sets among differentially expressed genes was tested using a Wilcoxon rank-sum test on the rank-ordered log2FC or p-values. For testing enrichment of the gene sets from Chen et al or Telese et al, 1-to-1 human orthologs were first obtained for the mouse genes using the orthogene package.

### Single-nucleus RNA sequencing data analysis

Raw sequencing reads in FASTQ format were processed using Cell Ranger (v5.0.0) for alignment to the hg38 genome and generation of gene expression matrices. Seurat (v4.0.3) in R was used for downstream analysis^88^. Doublets were removed using Scrublet^89^. In addition, nuclei with more than 2.5% mitochondrial RNA content, with less than 200 or more than 7,500 genes, or with UMI counts in the top 1% of all neurons were excluded. The remaining nuclei were integrated using Single Cell Transformation (SCT) within Seurat. Unsupervised clustering and identification of marker genes were performed on the integrated Seurat object. Specific cell types were manually annotated based on marker gene expression patterns. Cells belonging to individual cell types were extracted, generating separate Seurat objects for each type. Sub-clustering and marker gene identification were then performed on each individual cell type object within Seurat. Finally, pseudobulk analysis was conducted on each cell type object. Gene expression matrices were aggregated by sample and cluster, and differential expression analysis was performed using DESeq2^87^.

### Cell abundance statistical analysis

Statistical analysis of the differences in cell abundance for the different cell subpopulations identified with snRNA-seq was performed using scCODA, using all cell types as reference, and searching for clusters credible in more than half of time at an FDR < 0.05^22^.

### In-situ hybridization

#### Chromogenic ISH

In-situ hybridization was performed on formalin-fixed paraffin-embedded tissue sections using the RNAscope 2.5 HD Duplex chromogenic assay (ACDBio) following the manufacturer’s instructions. Briefly sections obtained from the NIH NeuroBioBank were baked for 1 hour at 60C. Sections were then deparaffinized, dehydrated in 100% ethanol and air-dried. Before hybridization, endogenous peroxidase activity was quenched by treating the sections for 10 minutes with hydrogen peroxide at room temperature. Sections were then boiled for 20 minutes in ACD target retrieval buffer. Last, they were incubated in ACD Protease Plus for 30 minutes at 40C. The hybridization of the C1 (RORB, ref 446061 or RELN, ref 413051) and C2 (GPC5 ref 521731-C2, BMPR1B, ref 468031-C2 or RELN, ref 413051-C2) probes, and detection was performed as recommended by the manufacturer except for the incubation times for Amp5, Amp6, Amp9 and Amp10 which were respectively 1 hour, 30 minutes, 1 hour and 30 minutes. Sections were counterstained with Gill’s hematoxylin 1, rinsed very quickly with tap water (the bluing step was omitted to preserve the green staining) and dried at 60C before being mounted in VectaMount mounting medium (ACD).

#### Multiplexed fluorescent ISH

For fluorescent ISH on human sections, we used the RNAscope multiplex fluorescent v2 kit, after photobleaching the sections to quench lipofuscin autofluorescence. For that purpose, we used the protocol described in Otero-Garcia et al., Neuron, 2022. Briefly, fresh frozen sections from C6-C8 were immersed in room temperature 10% buffered neutral formalin promptly after having been taken out of the – 80C freezer. They were fixed for 1 hour, washed twice in 1X PBS, dehydrated in increasing concentrations of ethanol and air-dried. They were then placed in photobleaching buffer (1X PBS with 0.1 mM Sodium Citrate pH = 7.2) for 72 hours at 4C under a 1,000 W full-spectrum LED lamp (around 3.5 inches away from the lamp), inside of a grow tent lined with high reflective mylar. Photobleaching buffer was replenished every day to compensate for evaporation. After photobleaching, the slides were dehydrated in 100% ethanol and air dried. Endogenous peroxidases were then quenched with hydrogen peroxide at room temperature, digested with protease IV for 30 minutes at room temperature, and then hybridize with the probe mixture as described in the multiplex fluorescent v2 kit. The probes were detected with either Opal 520, 570, 690 or 780.

### Yeast direct 1-by-1 interaction assay

The direct 1-by-1 interaction assay was performed by Hybrigenics Services SAS, Evry, France (http://www.hybrigenics-services.com).

The coding sequence of the APOER2 intracellular domain (NM_017522, Arg645 to Pro700) was cloned in frame with the LexA DNA binding domain (DBD) into pB27 as a C-terminal fusion to LexA (LexA-bait fusion). pB27 derives from the original pBTM116 vector^90^. The coding sequences of SNX17 (NM_014748.4), SNX27 (NM_001330723.2) and SNX31 (NM_152628.4) were cloned in frame with the Gal4 Activation Domain (AD) into plasmid pP7 (AD-prey fusion), derived from the original pGADGH^91^.

Bait and prey constructs were transformed into yeast haploid cells L40deltaGal4 (mata) and YHGX13 (Y187 ade2-101::loxP-kanMX-loxP, matα), respectively. The diploid yeast cells were generated by mating the two yeast strains^92^. These assays utilized the HIS3 reporter gene to monitor growth in histidine-deficient conditions. As negative controls, the bait plasmid was tested with empty prey vector (pP7) and all prey plasmids were tested with the empty bait vector (pB27). The interaction between SMAD and SMURF is used as positive control^93^.

All bait and prey combinations were tested for 1-by-1 interactions using three independent yeast clones on DO-2 and DO-3 selective media. The auxotrophy for tryptophan and leucine in DO-2 and DO-3 served as selective pressure for the maintenance of bait and prey plasmids. Clones maintained on DO-2 were then streaked onto DO-3 to assess direct 1-by-1 interactions. Only yeast clones exhibiting a bait/prey interaction were able to grow on DO-3.

### Immunofluorescence

Fresh frozen sections from the human hippocampal formation, including the EC cortex were post-fixed for 10 minutes in 4% PFA. They were then permeabilized in 0.1% Triton X-100, 2% normal goat or donkey serum in PBS for 30 min at room temperature. They were then stained with primary antibodies prepared in the same permeabilization buffer over night at 4°C. Primary antibodies were 6E10 (1:500, #803001, Biolegend), Reelin (1:50, #PA5-47537, Invitrogen), Dab1 (1:250, #LS-B9240, LSBio) and RBFOX1-Alexa594 (1:250, #862706, Biolegend). The day after samples were washed in PBS and incubated with secondary antibodies for 1h at room temperature. The secondary antibodies used were Goat anti-mouse IgG (H+L) Cross-adsorbed Alexa fluor 546 (1:500, #A-11003, ThermoFisher), Goat anti-rabbit IgG (H+L) Cross-adsorbed Alexa fluor 647 (1:500, #A-21245, ThermoFisher), Donkey anti-goat IgG (H+L) Cross-adsorbed Alexa fluor 488 (1:500, #A-11055, ThermoFisher), Donkey anti-rabbit IgG (H+L) Cross-adsorbed Alexa fluor 647 (1:500, #A32795, ThermoFisher) and Donkey anti-mouse IgG (H+L) Cross-adsorbed Alexa fluor 546 (1:500, #A10036, ThermoFisher). NFTs were stained with 1% Thioflavin S for 10 minutes at room temperature. After washing with PBS autofluorescence was quenched with Eliminator reagent (#2160, Millipore Sigma) following the manufacturer’s instructions and mounted with ProLong Gold Antifade Reagent (#P36980, ThermoFisher).

### Image acquisition and analysis

Immunofluorescence images were acquired in the Axio Scan Z1 automated slide scanner system (ZEISS) using the 20x objective. Brightfield whole-slide scans of chromogenic in-situ hybridization samples were acquired using a Vectra Polaris slide scanner (Akoya) using a 40x objective, and multiplexed fluorescent ISH with the 20x objective fluorescent slides were then unmixed using PhenoImager HT. Image analysis was performed with the QuPath software.

### Western Blotting analysis

Western blot analysis was performed on the supernatants obtained after tissue homogenization and centrifugation for nuclei isolation, that were further lysed with RIPA buffer supplemented with protease (#A32965, ThermoFisher), and phosphatase (#4906837001, PhoSTOP, Merck) inhibitors. Samples where then subjected to sodium dodecyl sulphate (SDS) polyacrylamide gel electrophoresis and transferred to a nitrocellulose membrane. Membranes were blocked in Intercept (TBS) blocking buffer (#927-60001, Li-Cor) 1h at room temperature and blotted with the primary antibodies overnight at 4 °C (mouse anti ApoE4 -4E4, #8941, Cell Signaling Technologies, 1:1,000), and goat anti-ApoE -AB947 (EMD Millipore, 1: 2,000). Membranes were then incubated with a Donkey anti-mouse IRDye 800CW and a Donkey anti-goat 680RD secondary antibodies (Li-cor, 1:10,000) for 1h at room temperature. Membranes were then stripped with NewBlot IR stripping buffer, blocked and blotted again with a mouse anti-actin primary antibody (NB600-501, Novus, 1:10,000) followed by a Donkey anti-mouse IRDye 800CW. Signal detection was performed with a Odyssey CLx scanning system (Li-Cor). Band intensity quantification was performed with the ImageStudio Lite software.

## Supporting information

Supplementary Data 1. Pseudo bulk DGE analysis and cluster markers

Supplementary Data 2. Pathway analysis for the different cell populations

Supplementary Data 3. FANS-seq results

## Abbreviations

AD=: Alzheimer’s Disease
ADN=: AD neuropathology
CGE=: caudal ganglionic eminence
CSF=: cerebrospinal fluid
DAM=: disease associated microglia
DEGs=: differentially expressed genes
DGE=: differential gene expression
EC=: entorhinal cortex
ECII=: entorhinal cortex layer II
FANS=: fluorescence activated nuclei sorting
GWAS=: genome wide association study
ISH=: in-situ hybridization
MGE=: medial ganglionic eminence
NFTs=: Neurofibrillary tangles
PFC=: prefrontal cortex
PET=: positron emission tomography.

## Acknowledgements

We thank C. Zhao, C. Lai and the Rockefeller University genomics resource center for the sequencing, T. Carroll and the Rockefeller University Bioinformatics Resource Center for their help with data analysis, S. Mazel, S. Semova, S. Han, S. Shalaby and the Rockefeller University Flow Cytometry Resource Center for nucleus sorting, A. O’Connell and H. Gertje and the Boston University Integrated Biomedical Imaging Services for their help with slide scan (with a shared equipment funded by NIH SIG grant S10OD030269 to N. Crossland), NIH NeuroBioBank (Harvard Brain Bank) for the human FFPE sections and the NBB (Netherlands Brain Bank) and Idibell biobanks for the fresh-frozen tissue samples, Dr. Eneritz Agirre for her support with snRNA-seq data analysis.

This work was supported by the European Union’s Horizon 2020 research and innovation program under the Marie Sklodowska-Curie grant agreement No 799638 (to P.R-R.); the Fisher Center for Alzheimer’s Disease Research (to J.P.R., P.R-R. and M.F.); Cure Alzheimer’s Fund (to J.P.R.); Alzheimerfonden (to P.R-R.); the Karolinska Institute fund for Doctoral education (KID) (to P.R-R.); Margaretha af Ugglas foundation (to S.M. and P.R-R.); The private initiative “Innovative ways to fight Alzheimer’s disease-Leif Lundblad family and others” (to P.R-R. and S.M.); Petrus and Augusta Hedlunds foundation (to P.R-R.); Gamla Tjänarinnor foundation (to P.R-R.,S.M. and C.T.); Gun and Bertil Stohnes foundation (to P.R-R., S.M. and C.T.); the National Institute on aging of the NIH (awards RF1 AG047779 and R21 AG085464) (to J.P.R.); Alzheimer’s Association (AARGD-22-932597) (to J.P.R.) and grants from the National Center for Advancing Translational Sciences (NCATS, National Institutes of Health (NIH) Clinical and Translational Science Award) #UL1TR001866 (CTSA) program and BU-CTSI #1UL1TR001430 (to J.P.R.). Its contents are solely the responsibility of the authors and do not necessarily represent the official views of the NIH.

## Author contribution

Authors contribution is assigned following CRediT (Contributor Roles Taxonomy) guidelines. Conceptualization and methodology, J.P.R. and P.R-R.; Formal Analysis, W.W., J.P.R. and P.R-R.; Investigation, P.R-R., C.T., I.F., A.M-M., K.J., V.A., H.A.K., E.M., X.L., A.M., and J.P.R.; Resources, N.V., S.M., P.R-R. and J.P.R.; Writing-Original Draft, J.P.R. and P.R-R.; Writing-Review & Editing, A.S-M., S.M., M.F., and A.M.; Visualization, P.R-R., W.W. and J.P.R.; Funding Acquisition, P.R-R and J.P.R.

## Data availability

The raw and processed snRNA-seq data is available at the gene expression omnibus (GEO) under accession code GSE287652 and RNA-seq data for sorted neuron nuclei is available under accession code GSE287651. Full raw slide-scanner images for immunofluorescence and in situ hybridization are available at Figshare https://doi.org/10.25452/figshare.plus.28255757.

## Extended data figures

**Extended Data Fig. 1. Sample cohort and pathology validation. a)** Immunofluorescence images of amyloid beta pathology (6E10, pink) and NFT (thioflavin S, green) in tissue sections from the different samples used for snRNAseq and FANS analysis. **b)** Western blot of ApoE4 (green) and total ApoE (red) in these same samples. Actin is used as loading control.

**Extended Data Fig. 2. Gene expression differences between control and ADN in different glia cells. a)** Volcano plot showing the differentially expressed genes between control and ADN in microglia and dysregulated pathways in these cells. **b)** Volcano plot showing the differentially expressed genes between control and ADN in astrocytes and dysregulated pathways in these cells. **c)** Volcano plot showing the differentially expressed genes between control and ADN in OPC and dysregulated pathways in these cells. **d)** Volcano plot showing the differentially expressed genes between control and ADN in oligodendrocytes and dysregulated pathways in these cells. **e)** Venn diagram of the overlap between differentially expressed genes in oligodendrocytes in ADN and plaque-induced genes (OLIGs) identified in the mouse.

**Extended Data Fig. 3. OPC and oligodendrocyte subpopulations and their abundance in control and ADN. a)** UMAP of the different OPC subpopulations and dot plot of the expression profiles for signature genes of prototypical subtypes. **b)** Upper panel: relative contribution of the different samples to a certain OPC subpopulation; lower panel: abundance of a certain OPC subtype in each sample. **c)** UMAP of the different oligodendrocyte subpopulations and dot plot of the expression profiles of prototypical subtypes signature genes. **d)** Upper panel: relative contribution of the different samples to a certain oligodendrocyte subpopulation; lower panel: abundance of a certain oligodendrocyte subtype in each sample.

**Extended Data Fig. 4. No differences in the abundance of different excitatory neuron subpopulations in early ADN. a)** UMAP of the integration of our dataset (blue) with Leng et al. (green) and the integration between control (blue) and ADN samples (red). **b)** Right panel: relative contribution of the different samples to a certain excitatory neuron subpopulation; right panel: abundance of a certain excitatory neuron subtype in each sample. **c)** Relative abundance of the RORB^+^ excitatory neuron subpopulation in our dataset and in Leng et al. **d)** In-situ hybridization images of *RELN* (red) and *RORB* (green) in the human EC. Scale bar 2 mm (a), 250 μm (b) and 20 μm (c-f).

**Extended Data Fig. 5. Gating strategy and neuron subpopulations sorted by FANS. a)** Gating strategy used for FANS of the different neuronal populations. **b**) Heatmap of the differentially expressed genes between NeuN^+^/RBFOX1^−^ and NeuN^+^/RBFOX1^+^ subpopulations and violin plots of their expression in the different inhibitory neuron subpopulations from snRNAseq. **c)** a. RBFOX1 signal intensity map of the different EC layers and RBFOX1fluorescence intensity quantifications per nucleus for each EC layers I-III and V-VI. b. RBFOX1 levels in Reelin positive neurons in the ECII. Scale bar 20 μm. **d)** Violin plots of *RBFOX1* and *LAMP5* expression in the different excitatory neuron subpopulations from snRNAseq. **e)** Wilcoxon test results for the enrichment of genes from the different excitatory neuron subpopulations in pvalue significant DEGs between the NeuN^+^/RBFOX1^−^ and NeuN^+^/RBFOX1^+^ (grey values) and for the enrichment of genes from the different excitatory neuron subpopulation based on Log2FC DEGs between NeuN^+^/RBFOX1^−^ and NeuN^+^/RBFOX1^+^ (blue and green values). **f)** ISH images of *PAX6, VIP, HTR3A, CBLN4, LAMP5* and *NPTX2* localization in the human EC. Scale bar 20 μm. **g)** ISH images showing the spatial localization of *LAMP5*+, *NPTX2*+, *LAMP5+/NPTX2+, CBLN4*+ and *LAMP5+/CBLN4*+ cells in the human EC. Scale bar 250 μm.

**Extended Data Fig. 6. Schematic representation of the molecular events taking place at early stages of AD neuropathology in the EC**. Under normal conditions, Reelin binds with ApoER2, signaling through its downstream effector Dab1. Our data indicate that Dab1 can shuttle between cytoplasm and nucleus, which possibly occurs through its interactions with the ApoER2 ICD, regulating transcription. During early stages of AD neuropathology, Reelin signaling is impaired and its effector Dab1, when present, localized in the cytosol in ECII neurons. This is accompanied by the upregulation of SNX31, an interactor of the ApoER2 ICD that might affect its recycling to the cell membrane. At the same time, the presence of amyloid plaques triggers the response of surrounding microglia. DAM microglia upregulate ApoE, which could further alter ApoER2 receptor availability at the cell surface. Concomitantly, pyramidal neurons in layers II/III of EC, downregulate NPTX2 and CBLN4, both essential regulator of the excitation inhibition balance of the region. This might be an effect of Reelin signaling impairment in neighboring stellate cells (NPTX2 is a Reelin transcriptional target). *Genes in loci or with mutations associated with resilience or increased AD risk by genetic studies. This image was generated with the help of Biorender.

