## Supplementary Table 1 for "Cell-type specific profiling of human entorhinal cortex at the onset of Alzheimer’s disease neuropathology"

| SAMPLE ID | BRAAK | CERAD | SEX | AGE | ApoE4 | Brain Bank | Sample type | Method |
|---|---|---|---|---|---|---|---|---|
| C1 | 0 | 0 | M | 54 | Yes | Idibell | FF/FFPE | snRNAseq/FANS/IF |
| C2 | 0 | 0 | F | 51 | No | Idibell | FF/FFPE | snRNAseq/FANS/IF |
| C3 | 0 | 0 | F | 60 | No | Idibell | FF/FFPE | snRNAseq/FANS/IF |
| C4 | 0 | 0 | M | 59 | No | Idibell | FF/FFPE | snRNAseq/FANS/IF |
| C5 | 0 | B | M | 71 | Yes | Idibell | FF/FFPE | snRNAseq/FANS/ |
| C6 | 0 | 0 | M | 64 | - | NBB | FF | ISH/IF |
| C7 | 0 | 0 | F | 64 | - | NBB | FF | ISH/IF |
| C8 | 0 | 0 | M | 68 | - | NBB | FF | ISH/IF |
| C9 | 0 | 0 | F | 55 | - | Harvard | FFPE | ISH |
| C10 | 0 | 0 | M | 36 | - | Harvard | FFPE | ISH |
| ADN1 | II | 0 | M | 70 | No | Idibell | FF/FFPE | snRNAseq/FANS/IF |
| ADN2 | II | B | F | 72 | No | Idibell | FF/FFPE | snRNAseq/FANS/IF |
| ADN3 | II | B | M | 71 | No | Idibell | FF/FFPE | snRNAseq/FANS/IF |
| ADN4 | II | B | M | 60 | No | Idibell | FF/FFPE | snRNAseq/FANS/IF |
| ADN5 | II | B | F | 78 | Yes | Idibell | FF/FFPE | snRNAseq/FANS |
| ADN6 | II | B | F | 79 | - | NBB | FF | ISH/IF |
| ADN7 | I | B | M | 66 | - | NBB | FF | ISH/IF |
| ADN8 | II | A | F | 67 | - | NBB | FF | ISH/IF |
