## Supplementary figures for "Cell-type specific profiling of human entorhinal cortex at the onset of Alzheimer’s disease neuropathology"

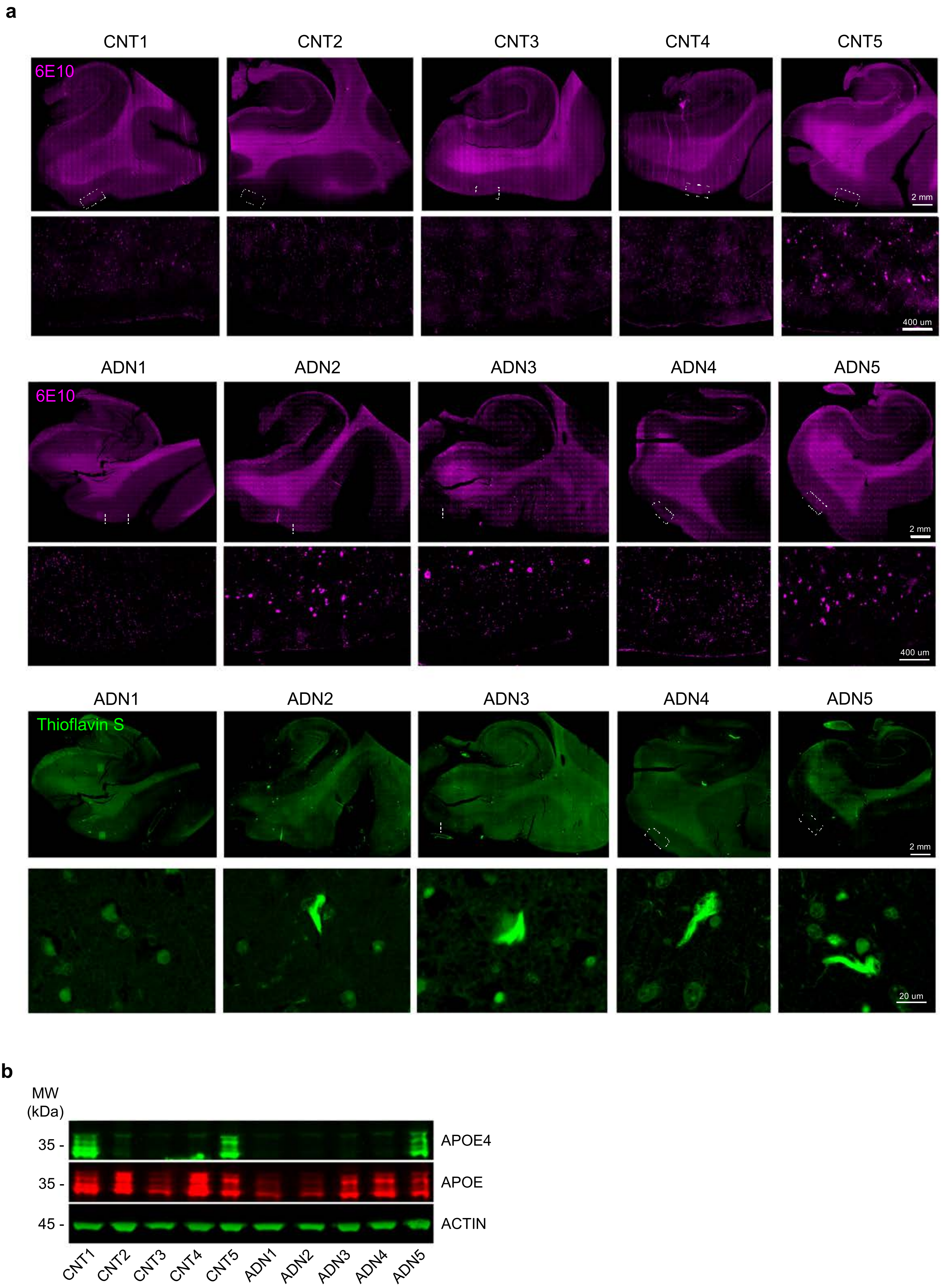
a
CNT1
CNT2
CNT3
CNT4
CNT5
6E10
2 mm
400 um
ADN1
ADN2
ADN3
ADN4
ADN5
6E10
2 mm
400 um
ADN1
ADN2
ADN3
ADN4
ADN5
Thioflavin S
2 mm
20 um
b
MW (kDa)
35 -
35 -
45 -
APOE4
APOE
ACTIN
CNT1
CNT2
CNT3
CNT4
CNT5
ADN1
ADN2
ADN3
ADN4
ADN5

Extended data Figure 2

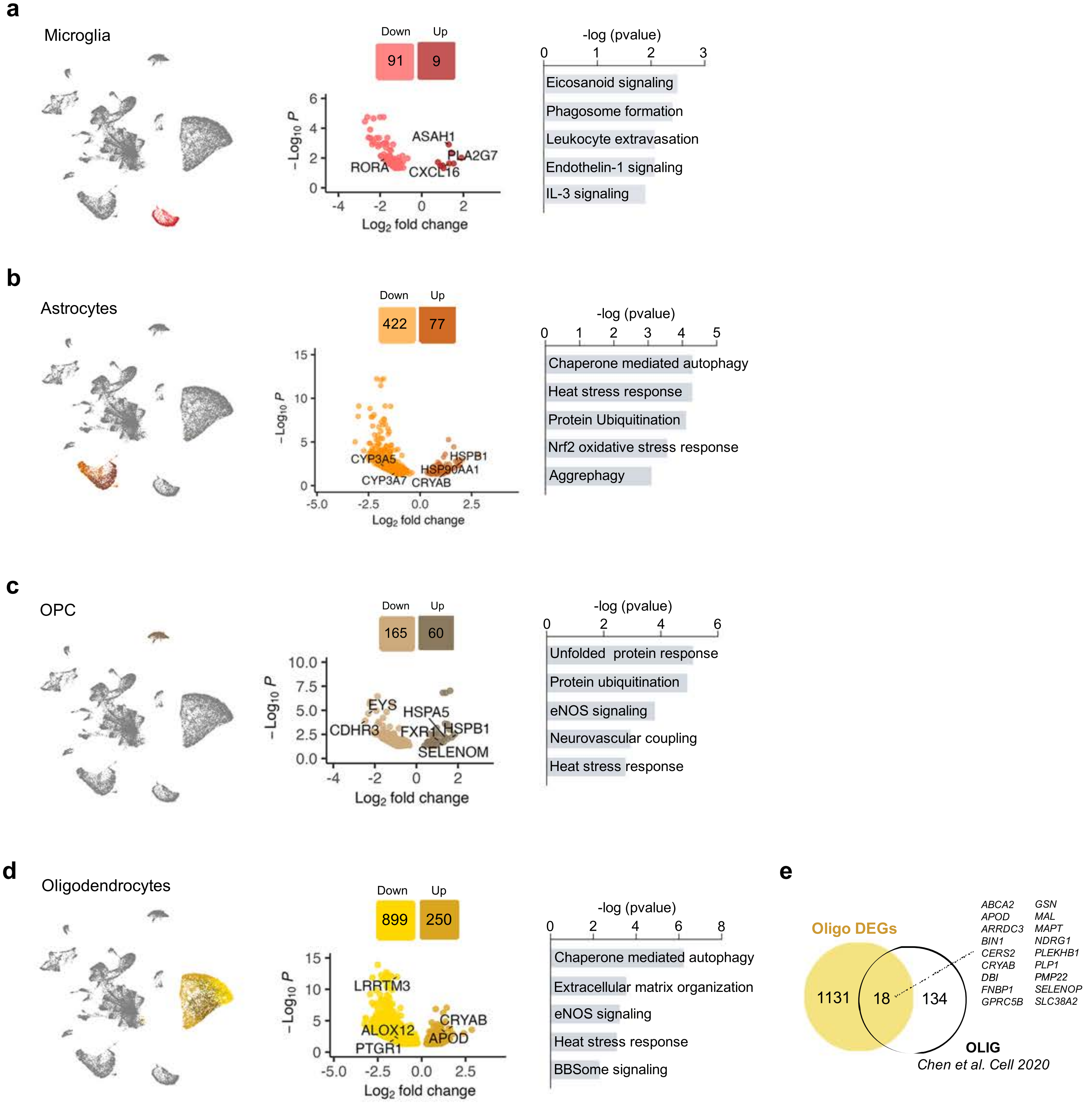

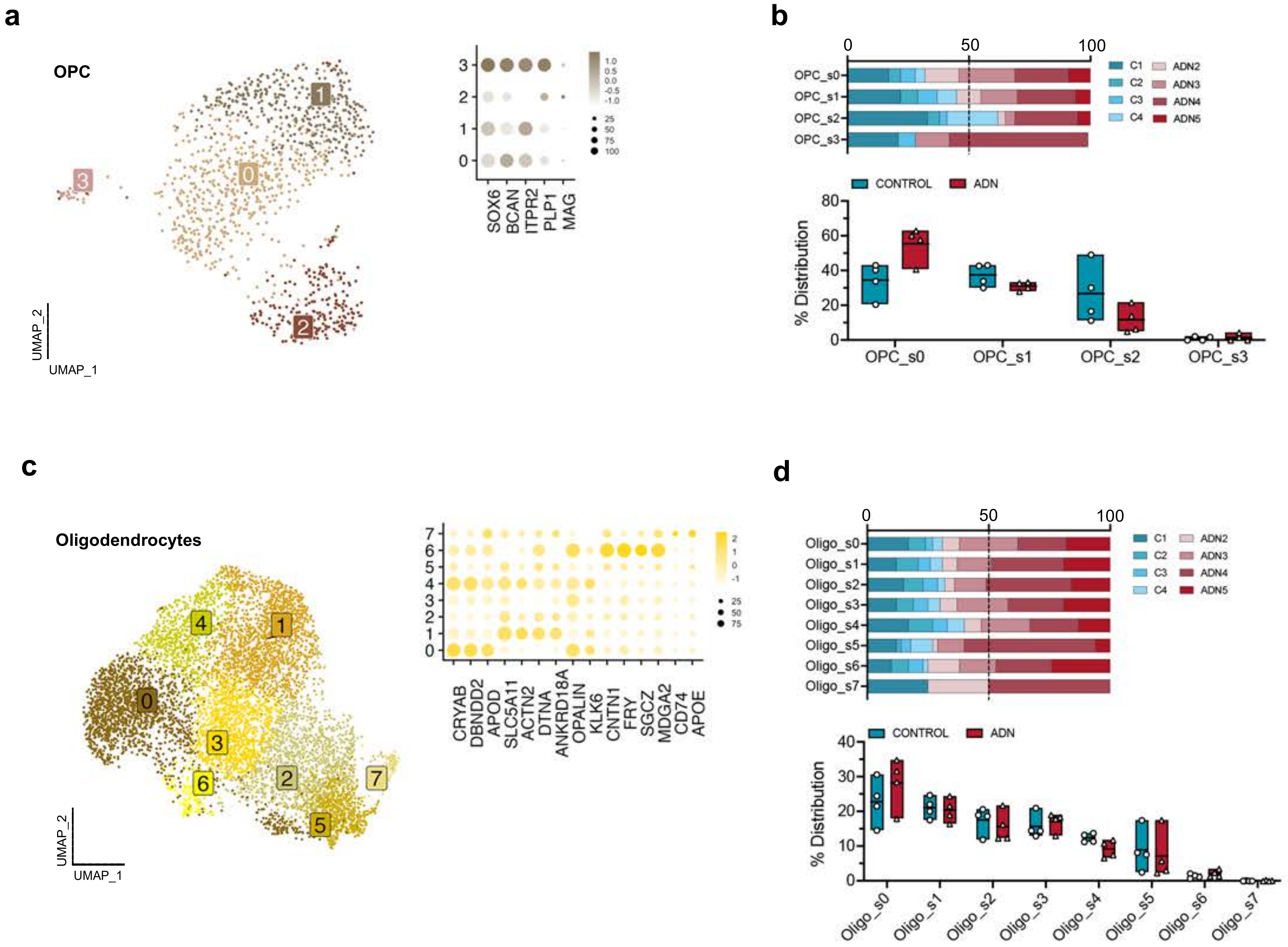
a
OPC
UMAP_2
UMAP_1
SOX6
BCAN
ITPR2
PLP1
MAG
b
OPC_s0
OPC_s1
OPC_s2
OPC_s3
C1
C2
C3
C4
ADN2
ADN3
ADN4
ADN5
CONTROL
ADN
% Distribution
c
Oligodendrocytes
CRYAB
DBNDD2
APOD
SLC5A11
ACTN2
DTNA
ANKRD18A
OPALIN
KLK6
CNTN1
FRY
SGCZ
MDGA2
CD74
APOE
d
Oligo_s0
Oligo_s1
Oligo_s2
Oligo_s3
Oligo_s4
Oligo_s5
Oligo_s6
Oligo_s7

Extended data Figure 4

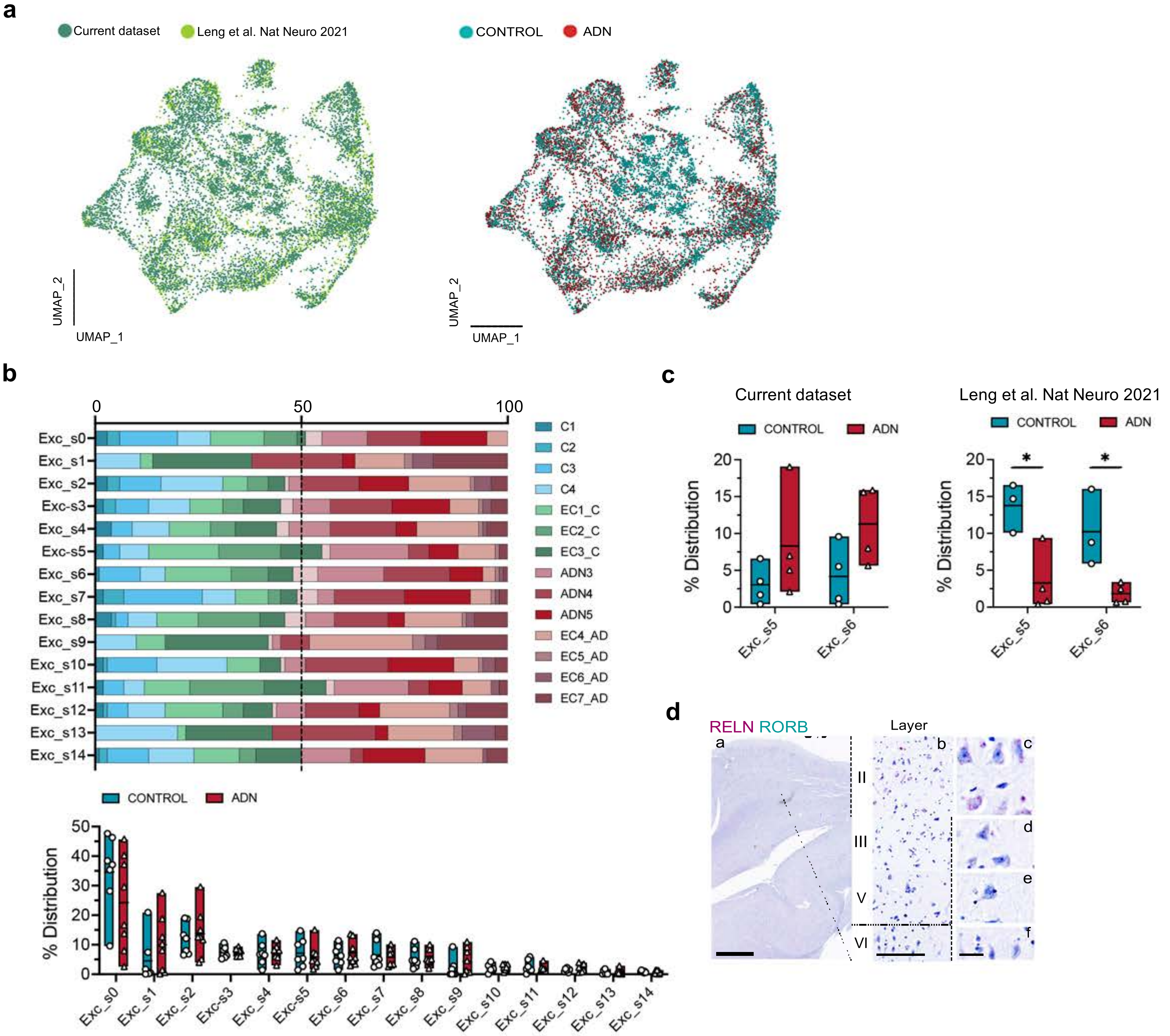

Extended data Figure 5

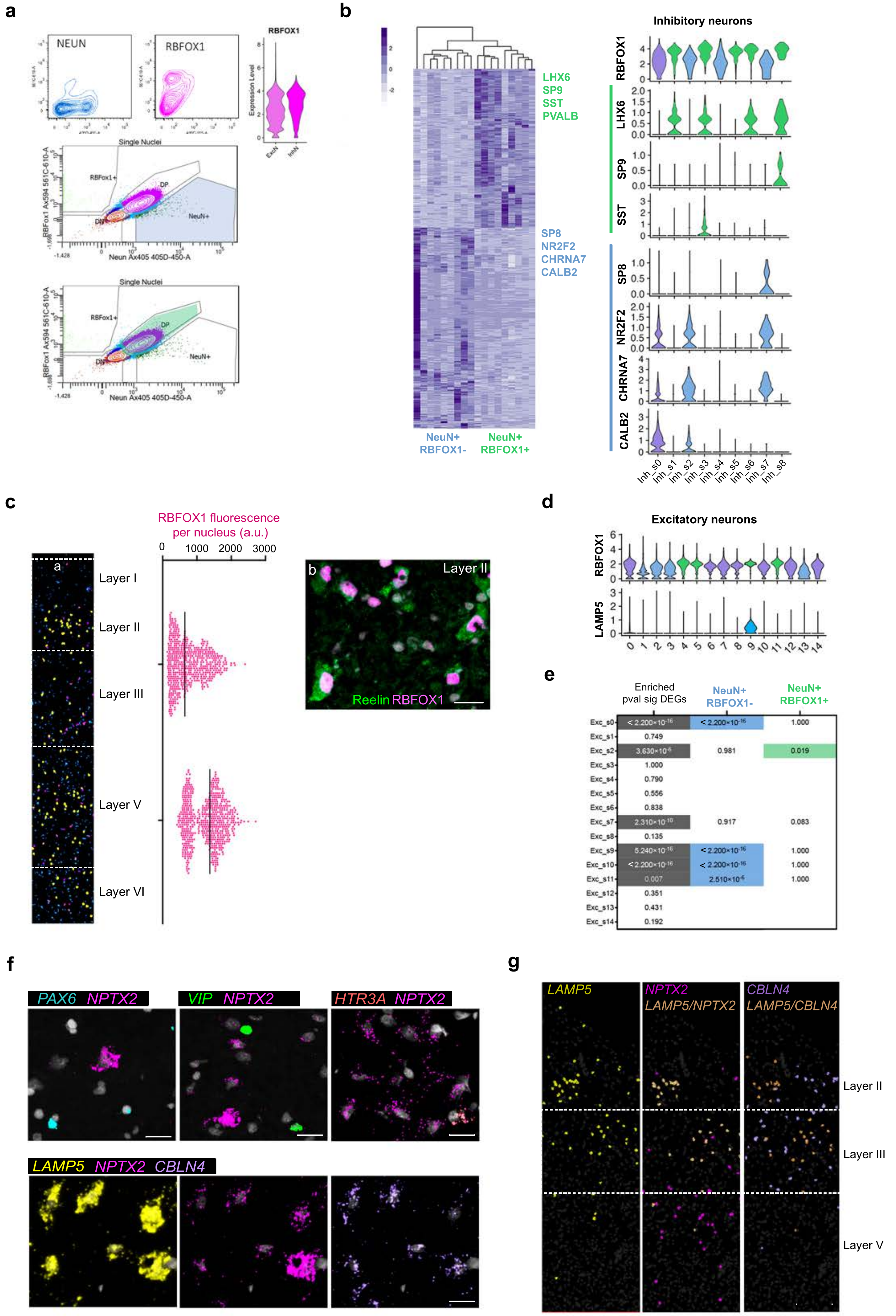


| | Enriched pval sig DEGs | NeuN+ RBFOX1- | NeuN+ RBFOX1+ |
|---|---|---|---|
| Exc_s0 | $< 2.200\times10^{-16}$ | $< 2.200\times10^{-16}$ | 1.000 |
| Exc_s1 | 0.749 | | |
| Exc_s2 | $3.630\times10^{-6}$ | 0.981 | 0.019 |
| Exc_s3 | 1.000 | | |
| Exc_s4 | 0.790 | | |
| Exc_s5 | 0.556 | | |
| Exc_s6 | 0.838 | | |
| Exc_s7 | $2.310\times10^{-10}$ | 0.917 | 0.083 |
| Exc_s8 | 0.135 | | |
| Exc_s9 | $5.240\times10^{-16}$ | $< 2.200\times10^{-16}$ | 1.000 |
| Exc_s10 | $< 2.200\times10^{-16}$ | $< 2.200\times10^{-16}$ | 1.000 |
| Exc_s11 | 0.007 | $2.510\times10^{-6}$ | 1.000 |
| Exc_s12 | 0.351 | | |
| Exc_s13 | 0.431 | | |
| Exc_s14 | 0.192 | | |

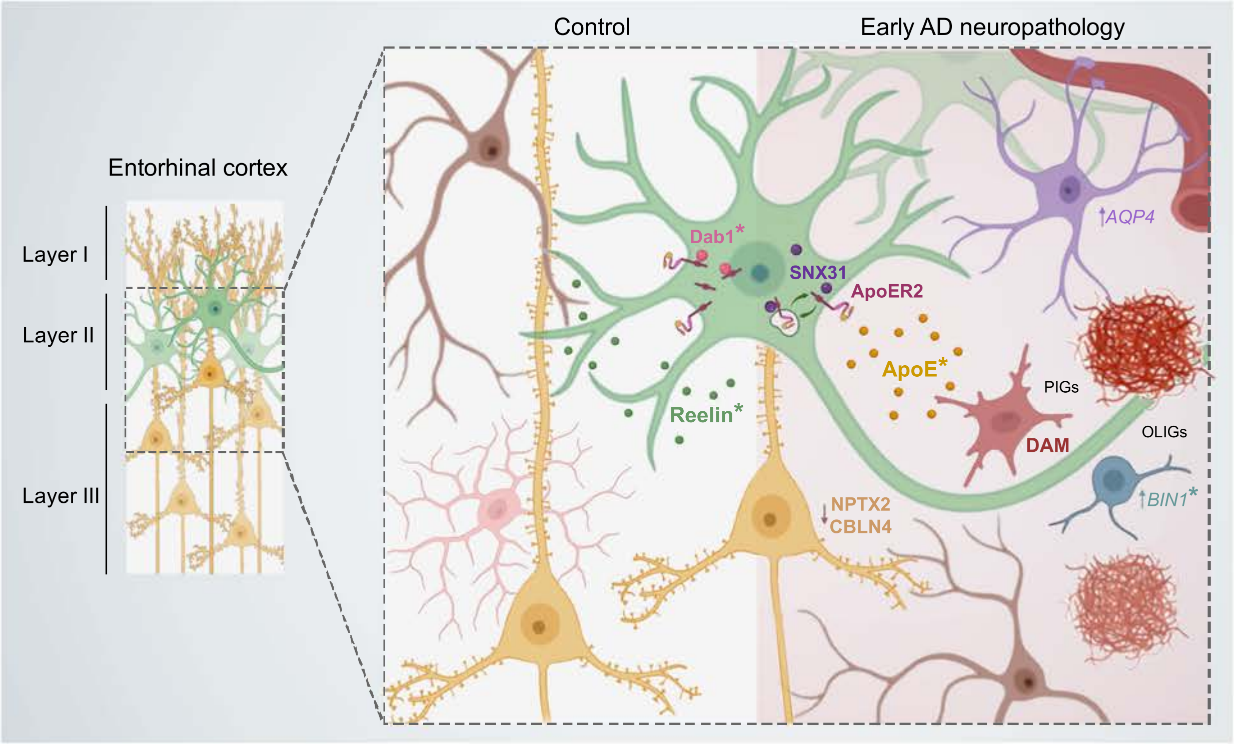
Control
Early AD neuropathology
Entorhinal cortex
Layer I
Layer II
Layer III
Dab1*
SNX31
ApoER2
ApoE*
Reelin*
AQP4
PIGs
DAM
OLIGs
BIN1*
NPTX2
CBLN4
